# IFNα differentially disrupts stimulus-secretion coupling in pancreatic alpha and beta cells

**DOI:** 10.64898/2026.10.20.760375

**Authors:** Atenea A. Perez-Serna, Daniel Guzman-Llorens, Fiona M. Gribble, Frank Reimann, Reinaldo S. dos Santos, Laura Marroqui

## Abstract

**Aims/hypothesis:** IFNα contributes to early islet inflammation and beta cell immunogenicity in type 1 diabetes, but its direct effects on endocrine cell function remain incompletely understood. We investigated whether acute IFNα exposure disrupts glucose-regulated insulin and glucagon secretion and whether the underlying alterations in stimulus-secretion coupling differ between beta and alpha cells.

**Methods:** Male and female mice received IFNα before an IPGTT, during which blood glucose and serum insulin and glucagon concentrations were measured. Hormone secretion and intracellular Ca^2+^ dynamics were assessed in isolated mouse islets and in MIN6, EndoC-βH1 and αTC1.9 cells following IFNα exposure. Sarco/endoplasmic reticulum Ca^2+^-ATPase 2 (SERCA2) protein levels and SERCA activity were measured in microsome-enriched fractions. Mitochondrial respiration was evaluated in intact and digitonin-permeabilised cells, including separate assessment of NADH-linked and succinate-linked pathways.

**Results:** Acute IFNα administration activated interferon signalling in isolated islets and altered glucose and hormone responses in vivo. IFNα increased glucose area under the curve in male mice but did not significantly affect glucose tolerance in female mice. The circulating insulin response to glucose was reduced in both sexes, while basal serum glucagon was reduced in female mice. In isolated islets, IFNα reduced glucose-stimulated insulin secretion in males and reduced 3-isobutyl-1-methylxanthine-potentiated insulin secretion in both sexes. IFNα also diminished glucose-mediated suppression of glucagon secretion in islets and attenuated the inhibitory effects of glucose and insulin in αTC1.9 cells. Intracellular Ca^2+^ signalling was impaired in primary islet cells and endocrine cell lines. IFNα did not significantly affect cell viability, reactive oxygen species production, proliferation or endocrine cell composition. IFNα reduced SERCA2 protein levels in MIN6 and EndoC-βH1 cells and impaired SERCA function in both models. By contrast, IFNα did not alter SERCA2 abundance or SERCA activity in αTC1.9 cells but reduced mitochondrial respiratory parameters. In permeabilised αTC1.9 cells, NADH-linked respiration was reduced, whereas succinate-linked respiration, mitochondrial mass and respiratory-chain complex abundance remained unchanged.

**Conclusions/interpretation:** Acute IFNα exposure disrupts endocrine cell stimulus-secretion coupling in the absence of cytotoxicity. Although intracellular Ca^2+^ signalling was impaired in both beta and alpha cells, the associated alterations were cell-type selective, involving reduced SERCA2 abundance and SERCA activity in beta cells and impaired NADH-linked mitochondrial respiration in alpha cells. These findings extend the pathogenic role of IFNα beyond immune activation and beta cell immunogenicity to include direct dysregulation of insulin and glucagon secretion.

## Introduction

Type 1 diabetes is characterised by the progressive loss of pancreatic beta cells through autoimmune mechanisms that develop before clinical diagnosis [1]. In addition to adaptive immune responses, innate inflammatory pathways contribute to disease initiation and progression. Among these, type I IFN, particularly IFNα, have emerged as key mediators of the early islet inflammatory response [2]. IFNα is expressed in human islets from individuals with type 1 diabetes [3, 4], and a type I IFN-inducible transcriptional signature has been detected before the onset of the disease [5, 6].

Studies in human islets and beta cell models have shown that IFNα induces sustained HLA class I overexpression, endoplasmic reticulum (ER) stress, and insulitis, three hallmarks of early type 1 diabetes [7–10]. Recent evidence further indicates that IFNα reshapes the beta cell immunopeptidome, promoting the presentation of alternative antigens and potentially increasing beta cell immunogenicity [11]. However, its impact on the cellular mechanisms governing hormone secretion remains incompletely understood.

Glucose-regulated hormone secretion relies on the coordinated interaction between cellular metabolism, intracellular Ca^2+^ signalling and exocytosis. In beta cells, nutrient metabolism drives ATP generation, membrane depolarisation and Ca^2+^ influx, ultimately promoting insulin secretion [12]. In alpha cells, glucagon secretion is regulated by intracellular Ca^2+^ dynamics, mitochondrial function and paracrine signals from neighbouring endocrine cells [13, 14]. Perturbations in mitochondrial metabolism or intracellular Ca^2+^ homeostasis can profoundly affect stimulus-secretion coupling and endocrine cell function [12, 13]. Recent work in human induced pluripotent stem cell-derived islet-like cells revealed that IFNα downregulates transcriptional pathways related to oxidative phosphorylation and mitochondrial function, suggesting that its effects may extend beyond its established immunomodulatory actions [15]. Nevertheless, the functional consequences of IFNα exposure on intracellular Ca^2+^ signalling and mitochondrial respiration in endocrine cells remain poorly understood.

Importantly, previous studies have focused largely on beta cells, whereas much less is known about the impact of IFNα on alpha cells. This is relevant because dysregulated glucagon secretion contributes to impaired glucose homeostasis in type 1 diabetes, and functional and metabolic abnormalities have been identified in alpha cells during early autoimmunity and established disease [16–19]. Whether IFNα affects stimulus-secretion coupling in beta cells and alpha cells through common or distinct mechanisms remains unknown.

Therefore, we investigated the effects of acute IFNα exposure on glucose-regulated insulin and glucagon secretion in vivo, in isolated islets and in endocrine cell models. We examined the cellular mechanisms underlying these effects, focusing on intracellular Ca^2+^ oscillations, sarco/endoplasmic reticulum Ca^2+^-ATPase 2 (SERCA2) protein levels and SERCA activity, and mitochondrial respiration in both beta cells and alpha cells.

## Methods

Additional methodological details are provided in the electronic supplementary material (ESM).

### Animals

Twelve-week-old male and female Glu-Venus mice expressing Venus under the control of the proglucagon promoter were used [20]. Mice were on a C57BL/6 background and were provided by Dr Fiona Gribble (University of Cambridge, UK). Twelve-week-old male and female C57BL/6 mice not carrying the Glu-Venus transgene were also used in some experiments. No a priori statistical calculation was performed to determine the sample size for the animal experiments, and animals were not randomly allocated to experimental groups. Investigators were not blinded to group allocation during the experiments or when assessing outcomes. Experimental procedures complied with the Spanish Royal Decree 1201/2005 and the European Community Council Directive 2010/63/EU. All protocols were reviewed and authorized by the Ethics Committee of Universidad Miguel Hernández de Elche (Alicante, Spain) (protocol ID: UMH.IDI.LME.240119).

### IFNα challenge and IPGTT

Male and female mice received intraperitoneal injections of saline or IFNα (10,000 IU) at the beginning of an overnight fast and again 2 h before the IPGTT. Animals remained fasted throughout the treatment period. Following intraperitoneal administration of D-glucose (2 g/kg body weight), blood glucose concentrations were measured in tail vein blood at 0, 15 and 120 min using an Accu-Chek glucometer (Roche, Basel, Switzerland). Blood samples (50 μl) were collected at 0 and 120 min for determination of serum insulin and glucagon concentrations using ELISA kits (Mercodia, Uppsala, Sweden).

### Islet isolation and cell culture

Mouse islets were isolated using collagenase (Sigma-Aldrich, St Louis, MO, USA) and, when required, dispersed with trypsin-EDTA and plated on poly-L-lysine-coated plates [21, 22]. Whole islets recovered for 2 h, whereas dispersed cells were cultured for 48 h before experimental treatments.

MIN6, αTC1.9 and EndoC-βH1 cells were cultured as previously described [23–26] and routinely tested negative for *Mycoplasma* contamination.

### IFNα treatment

MIN6 cells, αTC1.9 cells and isolated mouse islets were treated with recombinant universal IFNα (1000 U/ml; PBL Assay Science, Piscataway, NJ, USA), whereas EndoC-βH1 cells were treated with recombinant human IFNα (1000 U/ml; R&D Systems, Minneapolis, MN, USA). For EndoC-βH1 cell treatments, culture medium was supplemented with 2% FBS. Cells and islets were exposed to IFNα for 24 h unless otherwise indicated.

### Intracellular Ca^2+^ recordings

Intracellular Ca^2+^ dynamics were assessed in intact islets and cultured cells using fluorescence imaging [27–30]. EndoC-βH1, MIN6 and αTC1.9 cells and whole islets were loaded with 2 μmol/l Fura-2AM, whereas primary alpha cells within intact islets were loaded with 2 μmol/l Fluo-4AM. After a basal period under perfusion with Krebs-Ringer bicarbonate buffer, cells were challenged with stimulatory glucose concentrations according to the experimental model, followed by depolarising KCl or adrenaline stimulation to assess glucose-independent Ca^2+^ responses. Fluorescence was recorded every 2 s.

For Fura-2 experiments, intracellular Ca^2+^ responses were quantified as changes in the 340/380 fluorescence ratio. For Fluo-4 experiments, responses were expressed as F/F_0_. Depending on the experimental model, Ca^2+^ activity was analysed as AUC, peak amplitude, oscillation frequency or inhibition/stimulation indices. See ESM Methods for further details.

### Hormone secretion

Insulin and glucagon secretion and content were assessed in isolated mouse islets, MIN6 cells and αTC1.9 cells using static incubation assays [23, 24, 31]. Following preincubation under basal glucose conditions, islets and cells were exposed to model-specific glucose concentrations and secretagogues. Insulin and glucagon concentrations in supernatants were quantified by ELISA. Cellular hormone content was determined from HCl-ethanol extracts. See ESM Methods for further details.

### Subcellular fractionation

Microsome-enriched fractions were obtained by differential centrifugation of cellular homogenates using an adapted protocol [32]. Briefly, cells were homogenised in ice-cold sucrose-based buffer and subjected to sequential centrifugation. The resulting microsomal pellet was collected and stored at −80°C until use. See ESM Methods for further details.

### Immunoblotting

Western blot analysis was performed on cell lysates or subcellular fractions as previously described [33, 34]. Proteins were separated by SDS-PAGE, transferred to PVDF membranes, and probed with specific primary and HRP-conjugated secondary antibodies (ESM Table 1).

### SERCA activity

SERCA activity was assessed in microsome-enriched fractions using a colorimetric ATPase/GTPase activity assay kit (Sigma-Aldrich). SERCA-specific activity was determined as cyclopiazonic acid-sensitive ATPase activity and normalised to protein content. Absorbance was measured using an iMark Microplate Reader (Bio-Rad, Hercules, CA, USA). See ESM Methods for further details.

### High-resolution respirometry

Mitochondrial respiration was measured in intact and digitonin-permeabilised MIN6, EndoC-βH1 and αTC1.9 cells using high-resolution respirometry (Oxygraph-2k, OROBOROS Instruments, Innsbruck, Austria). Intact-cell respiratory parameters and NADH- and succinate-linked respiration in permeabilised cells were assessed using modified SUIT-003 and SUIT-008 protocols, respectively. Oxygen consumption rates (OCR) were corrected for residual oxygen consumption and analysed using O2k-Software DatLab 7.4. See ESM Methods for further details.

### qRT-PCR

RNA was extracted using the NucleoSpin RNA Kit (Macherey-Nagel, Düren, Germany) following the manufacturer’s instructions. cDNA synthesis and quantitative PCR were performed as described [29]. Values were analysed with CFX Manager Version 1.6 (Bio-Rad) and expressed relative to control values using the 2^-ΔCt^ method. *Hprt1* was used as housekeeping gene. The primers are listed in ESM Table 2.

### Statistical analysis

Data are presented as mean ± SEM, unless otherwise stated. The experimental unit and exact sample size are defined in each figure legend. Comparisons between two groups were performed using paired or unpaired two-tailed Student’s *t* tests, as determined by the experimental design. Comparisons involving treatment and stimulus or glucose concentration were performed using two-way ANOVA or mixed-effects analysis, followed by Sidak’s or Tukey’s multiple-comparisons test, as indicated. Data distribution was assessed using the Shapiro-Wilk test, and non-normally distributed data were analysed using the corresponding non-parametric tests. Differences were considered statistically significant at *p*≤0.05. Statistical analyses were performed with GraphPad Prism 8 software (GraphPad Software, San Diego, CA, USA).

## Results

### Acute IFNα exposure activates interferon signalling in islets and alters glucose-regulated hormone secretion in vivo

To assess the acute effects of IFNα on glucose homeostasis and islet hormone secretion in male and female mice, animals were challenged with IFNα prior to an IPGTT (Fig. 1a). Increased expression of the canonical interferon-stimulated genes *Mx1* and *Cxcl10* confirmed activation of interferon signalling in isolated male (Fig. 1b, c) and female islets (Fig. 1d, e).

**Fig. 1.**
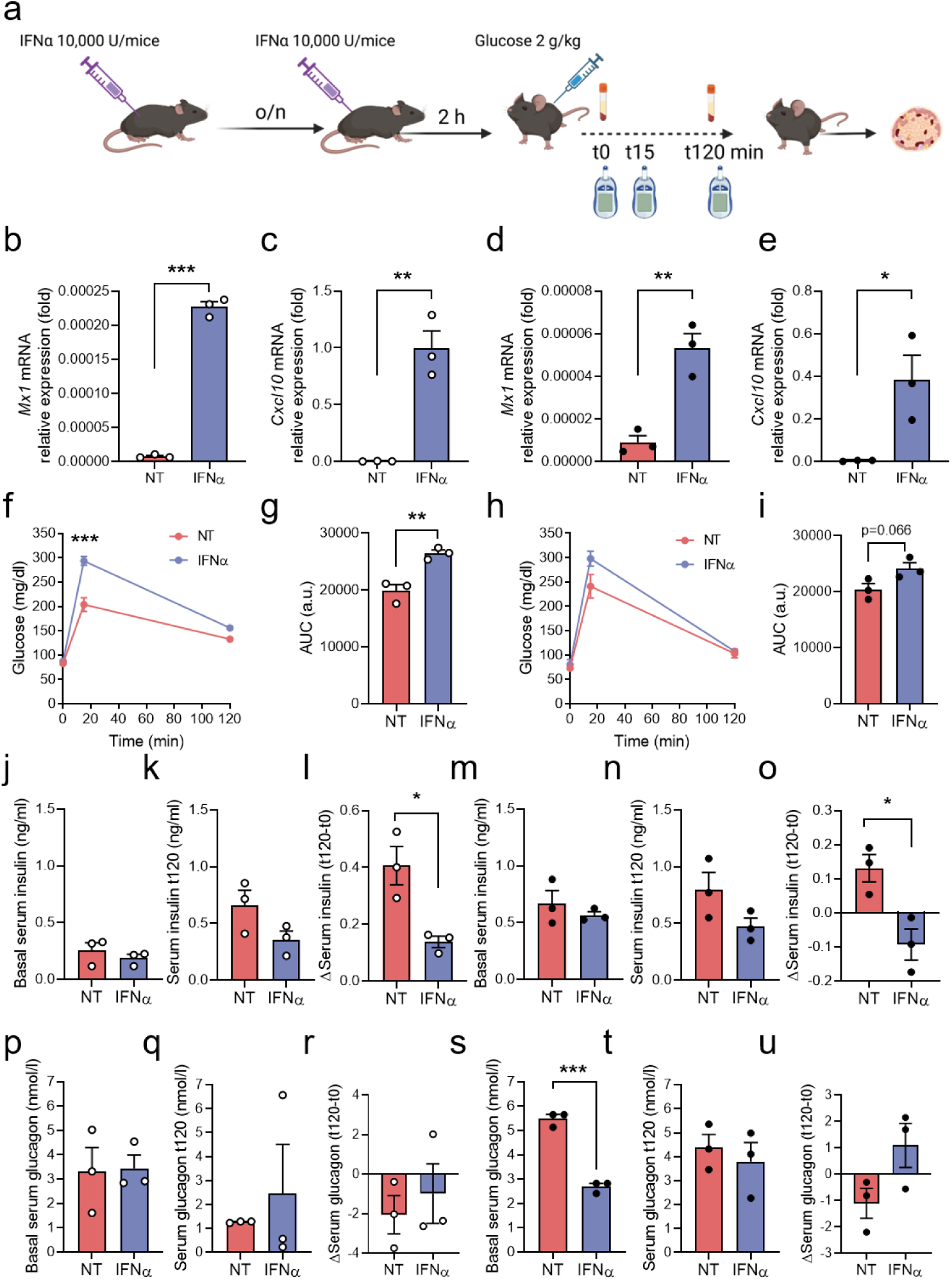
Acute in vivo exposure to IFNα impairs glucose tolerance in mice. (**a**) Experimental protocol. Male mice (open circles) and female mice (closed circles) received intraperitoneal injections of saline (NT; red) or IFNα (10,000 IU; blue) at the beginning of an overnight fast and again 2 h before the IPGTT, followed by intraperitoneal administration of glucose (2 g/kg body weight). Blood glucose was measured at 0, 15 and 120 min, and blood samples were collected at 0 and 120 min for measurement of serum insulin and glucagon concentrations. At 120 min, mice were euthanised and pancreatic islets isolated for gene expression analysis. (**b-e**) mRNA expression of *Mx1* (**b, d**) and *Cxcl10* (**c, e**) was analysed by qRT-PCR and normalised to *Hprt1*. (**f, h**) Blood glucose concentrations and (**g, i**) glucose AUC during the IPGTT. Serum insulin concentrations at 0 min (**j, m**) and 120 min (**k, n**), and changes in circulating insulin during the IPGTT (**l, o**). Serum glucagon concentrations at 0 min (**p, s**) and 120 min (**q, t**), and changes in circulating glucagon during the IPGTT (**r, u**). Serum insulin and glucagon were measured by ELISA. Results are means ± SEM of three male and three female mice per condition. \**p*≤0.05, \*\**p*≤0.01 and \*\*\**p*≤0.001 vs NT; unpaired Student’s *t* test or two-way ANOVA followed by Sidak’s test for multiple comparisons, as appropriate.

Blood glucose measurements during the IPGTT revealed different response patterns in male and female mice. IFNα increased glycaemia during the IPGTT in male mice (Fig. 1f), resulting in a 34% higher glucose AUC than in untreated animals (Fig. 1g). In female mice, despite higher glucose concentrations in untreated animals at 15 min, IFNα did not significantly alter glucose AUC (Fig. 1h,i). Of note, no significant difference in glucose AUC was detected between IFNα-treated male and female mice.

To investigate whether these changes were associated with altered endocrine responses, serum insulin and glucagon concentrations were measured during the IPGTT. In male mice, IFNα did not significantly affect serum insulin concentrations at baseline or 120 min (Fig. 1j, k). However, the change in circulating insulin between baseline and 120 min was smaller following IFNα treatment (Fig. 1l). Similarly, serum insulin concentrations did not differ significantly between groups in female mice, although concentrations at 120 min were numerically lower following IFNα treatment (Fig. 1m,n). The change in insulin between baseline and 120 min was also reduced in IFNα-treated female mice (Fig. 1o). Serum glucagon concentrations were unchanged in male mice, and the change between baseline and 120 min did not differ significantly between groups (Fig. 1p-r). In female mice, basal serum glucagon concentrations were lower following IFNα treatment than in untreated controls (Fig. 1s), whereas concentrations at 120 min and the change between baseline and 120 min were not significantly different between groups (Fig. 1t,u). These findings indicate that acute IFNα exposure alters glucose-regulated hormone responses in vivo, prompting us to investigate its direct effects on islet endocrine function.

### IFNα impairs insulin and glucagon secretion in islets and cell lines

To determine whether IFNα directly affects endocrine cell function, insulin and glucagon secretion were assessed in mouse islets, MIN6 cells and αTC1.9 cells (Fig. 2). In male islets, IFNα decreased glucose-stimulated insulin secretion (GSIS) by 25% (Fig. 2a), while no changes were observed in islets from female mice (Fig. 2c). Nevertheless, IFNα reduced 3-isobutyl-1-methylxanthine (IBMX)-induced insulin secretion by 20-30% in islets from both sexes (Fig. 2b, d).

**Fig. 2.**
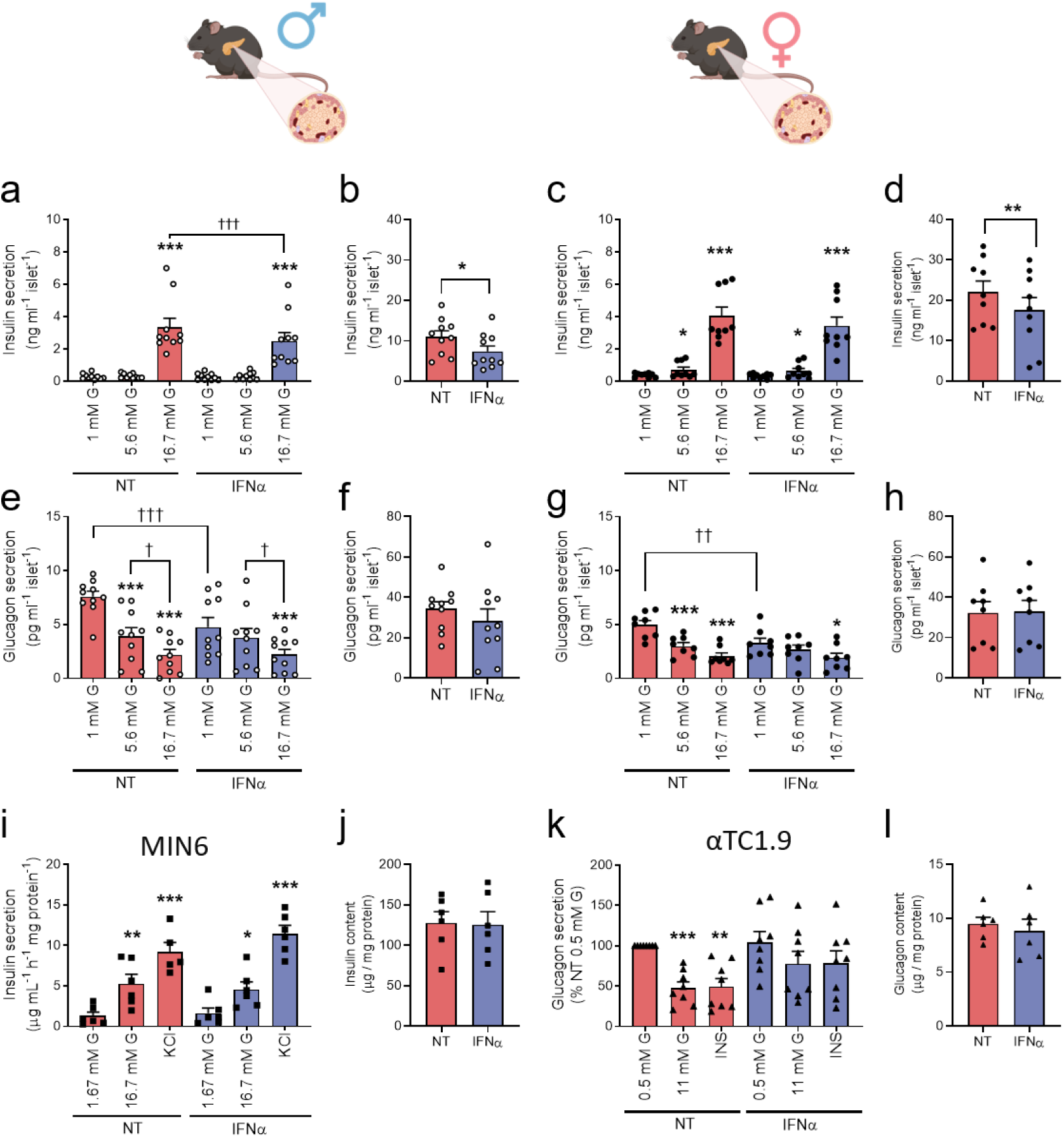
IFNα impairs glucose-regulated insulin and glucagon secretion. Islets from male (open circles) and female (closed circles) GCG-Venus mice were left untreated (NT; red) or treated with IFNα (1000 U/ml; blue) for 24 h. Insulin (**a, c**) and glucagon (**e, g**) secretion in response to 1, 5.6 and 16.7 mmol/l glucose were measured simultaneously in the same islets. IBMX-potentiated insulin (**b, d**) and glucagon (**f, h**) secretion were also assessed. MIN6 cells (**i, j**; squares) and αTC1.9 cells (**k, l**; triangles) were left untreated (NT, red) or treated with IFNα (1000 U/ml, blue) for 24 h. Insulin secretion (**i**) and content (**j**) in MIN6 cells. Glucagon secretion (**k**) and content (**l**) in αTC1.9 cells. Insulin and glucagon were measured by ELISA. Results are means ± SEM of 6-10 independent experiments. For (**a, c, e, g, i, k**), \**p*≤0.05, \*\**p*≤0.01 and \*\*\**p*≤0.001 vs low glucose (i.e., 1, 1.67 or 0.5 mmol/l); ^†^*p*≤0.05, ^††^*p*≤0.01 and ^†††^*p*≤0.001 as indicated by bars; two-way ANOVA followed by Sidak’s test for multiple comparisons. For (**b, d, f, h, j, l**), \**p*≤0.05 and \*\**p*≤0.01 vs NT; paired Student’s *t* test.

In untreated islets from both sexes, glucagon secretion decreased relative to basal glucose conditions (1 mmol/l glucose) by 50% and 70% at 5.6 mmol/l and 16.7 mmol/l glucose, respectively (Fig. 2e, g). Conversely, this glucose-mediated suppression was attenuated by IFNα in both sexes, with reductions of only 20% and 55-60% at 5.6 and 16.7 mmol/l glucose, respectively (Fig. 2e, g). IFNα also reduced glucagon secretion at 1 mmol/l glucose. By contrast, IBMX-potentiated glucagon secretion was not significantly affected by IFNα in islets from either sex (Fig. 2f, h).

To determine whether the effects of IFNα observed in isolated islets reflected direct actions on endocrine cells, hormone secretion and content were assessed in MIN6 and αTC1.9 cells (Fig. 2i-l). In contrast to male islets, IFNα did not alter GSIS in MIN6 cells, nor did it affect insulin secretion in response to KCl (Fig. 2i). In αTC1.9 cells, IFNα did not alter glucagon secretion under stimulatory low-glucose conditions (0.5 mmol/l glucose) (Fig. 2l) but attenuated the inhibitory effects of 11 mmol/l glucose and insulin (Fig. 2k). Whereas glucagon secretion was suppressed by approximately 50% in untreated cells, the inhibitory effects of glucose and insulin were reduced to approximately 20% following IFNα exposure (Fig. 2k). Insulin and glucagon content remained unchanged in IFNα-treated MIN6 and αTC1.9 cells, respectively (Fig. 2j, l).

The different effects in islets and cell lines suggest that intra-islet cell interactions may contribute to the impairment of insulin secretion observed ex vivo. In contrast, the αTC1.9 phenotype supports a direct effect of IFNα on alpha cell function.

### IFNα does not affect cell viability, reactive oxygen species production, proliferation or endocrine cell composition

As IFNα is known to induce inflammatory and ER stress responses [7, 10], we assessed whether the observed secretory defects were associated with alterations in cell viability, intracellular reactive oxygen species (ROS) production, proliferation or endocrine cell composition. IFNα did not affect cell viability in MIN6, αTC1.9 and EndoC-βH1 cells, nor in dispersed islet cells from male and female mice (ESM Fig. 1a-c, h, i). Likewise, no changes in ROS production were detected in MIN6, αTC1.9 or EndoC-βH1 cells following IFNα treatment (ESM Fig. 1e-g). Cell proliferation and the proportions of insulin-positive and glucagon-positive cells in dispersed islets were also unchanged (ESM Fig. 2).

### IFNα impairs intracellular Ca^2+^ oscillations in islet cells

As the secretory defects induced by IFNα could not be explained by alterations in cell viability, oxidative stress or endocrine cell composition, we next investigated whether IFNα affects intracellular Ca^2+^ oscillations, a central regulator of insulin and glucagon secretion. The analysed Ca^2+^ parameters are summarised in ESM Fig. 3.

**Fig. 3.**
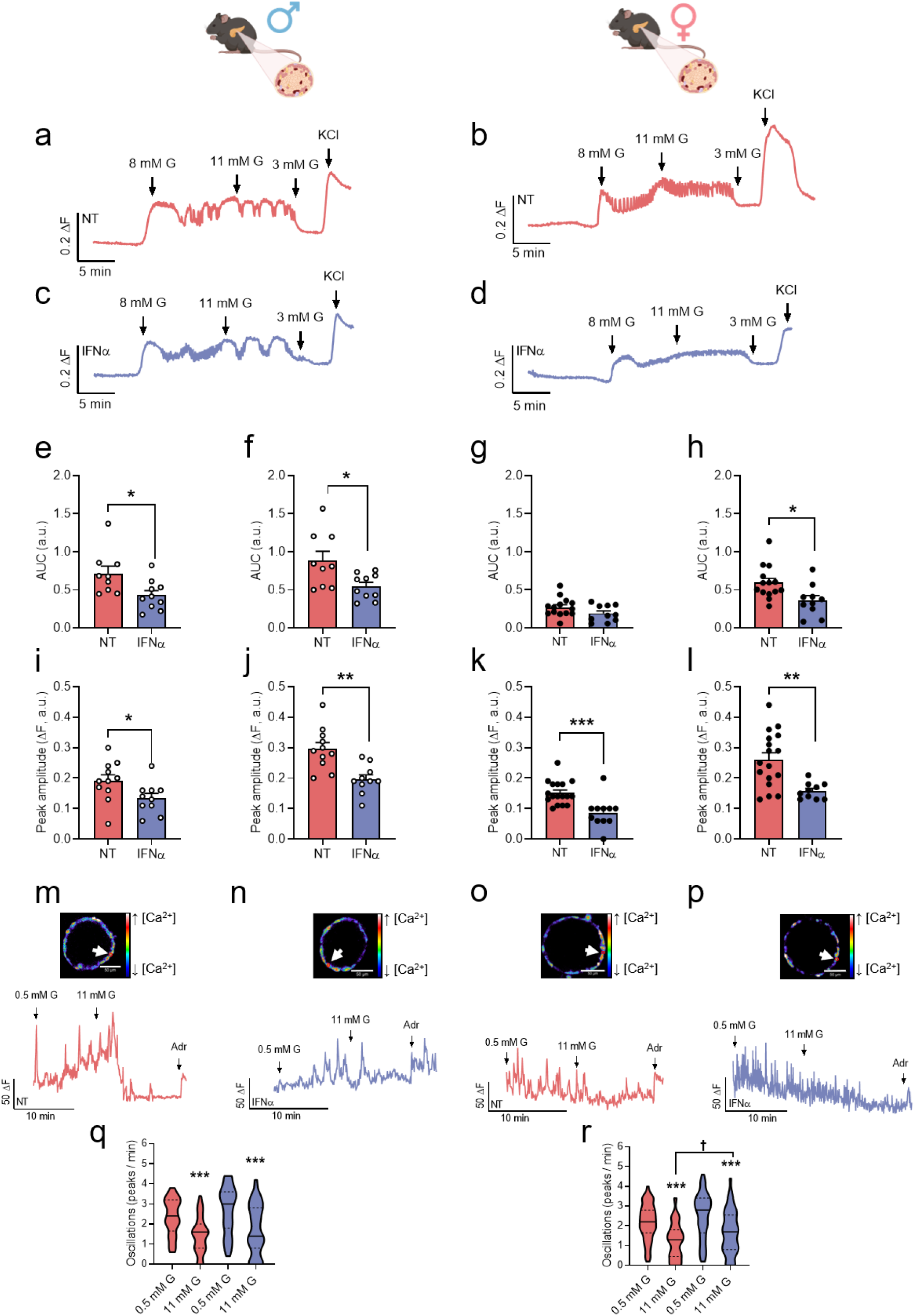
IFNα impairs intracellular Ca^2+^ oscillations in pancreatic islets. Islets from male (open circles) and female (closed circles) GCG-Venus mice were left untreated (NT; red) or treated with IFNα (1000 U/ml; blue) for 24 h. (**a-d**) Representative recordings of intracellular Ca^2+^ from individual intact islets exposed sequentially to 8, 11 and 3 mmol/l glucose, and 30 mmol/l KCl. Islets were left untreated (**a, b**; red) or treated with IFNα (**c, d**; blue). (**e-h**) Quantification of AUC in intracellular Ca^2+^ changes in response to 8 mmol/l glucose (**e, g**) and 11 mmol/l glucose (**f, h**). (**i-l**) Peak amplitude of the initial Ca^2+^ response to 8 mmol/l glucose (**i, k**) and KCl (**j, l**). Results are means ± SEM from 10-11 registers (NT: *n*=11 islets; IFNα: *n*=10 islets) from four male mice (**e, f, i, j**) and 10-17 registers (NT: *n*=14-17 islets; IFNα: *n*=10 islets) from five female mice (**g, h, k, l**). Each dot represents an individual register. \**p*≤0.05, \*\**p*≤0.01 and \*\*\**p*≤0.001 vs NT; unpaired Student’s *t* test. (**m-p**) Representative recordings of intracellular Ca^2+^ from individual primary alpha cells exposed sequentially to 0.5 and 11 mmol/l glucose, and 5 μmol/l adrenaline (Adr). Cells were analysed in untreated (**m, o**; red) or IFNα-treated islets (**n, p**; blue) from male (**m, n**) and female (**o, p**) mice. Individual representative cells are indicated by white arrows. Scale bar, 50 μm. (**q, r**) Ca^2+^ oscillation frequency in alpha cells from male (**q**) and female (**r**) mice. Violin plots show the distribution, median (solid lines) and quartiles (dashed lines) of recordings from individual alpha cells. (**q**) 59-60 registers (NT: *n*=18 islets; IFNα: *n*=16 islets) from eight male mice. (**r**) 52-57 registers (NT: *n*=9 islets; IFNα: *n*=10 islets) from four female mice. \*\*\**p*≤0.001 vs 0.5 mmol/l glucose within each condition; ^†^*p*≤0.05 as indicated by bars; two-way ANOVA followed by Sidak’s test for multiple comparisons. G: Glucose.

IFNα reduced the overall Ca^2+^ response to glucose in islets from both sexes, as shown by the representative traces (Fig. 3a-d). In male islets, AUC was reduced by approximately 40% at both 8 mmol/l and 11 mmol/l glucose (Fig. 3e, f), while female islets showed a 31% reduction at 11 mmol/l glucose (Fig. 3h). IFNα also decreased peak Ca^2+^ amplitude in response to 8 mmol/l glucose by 28% in male and 38% in female islets (Fig. 3i, k). Moreover, depolarisation-induced Ca^2+^ responses was impaired, as KCl-evoked peak amplitude was similarly reduced by 34% and 40% in males and females, respectively (Fig. 3j, l).

Ca^2+^ dynamics were analysed in primary alpha cells identified by their characteristic glucose- and adrenaline-responsive Ca^2+^ oscillatory profile (Fig. 3m-p). In contrast to beta cells, IFNα did not affect Ca^2+^ oscillations in males (Fig. 3q) but reduced the inhibitory effect of 11 mmol/l glucose by 33% in females (Fig. 3r).

Collectively, these findings indicate that IFNα impaired beta cell Ca^2+^ oscillations in both sexes, while its effects on primary alpha cells were predominantly observed in females.

### IFNα impairs intracellular Ca^2+^ oscillations in beta and alpha cell models

To assess direct effects on endocrine cells, intracellular Ca^2+^ dynamics were analysed in MIN6, EndoC-βH1 and αTC1.9 cells following IFNα exposure (Fig. 4).

**Fig. 4.**
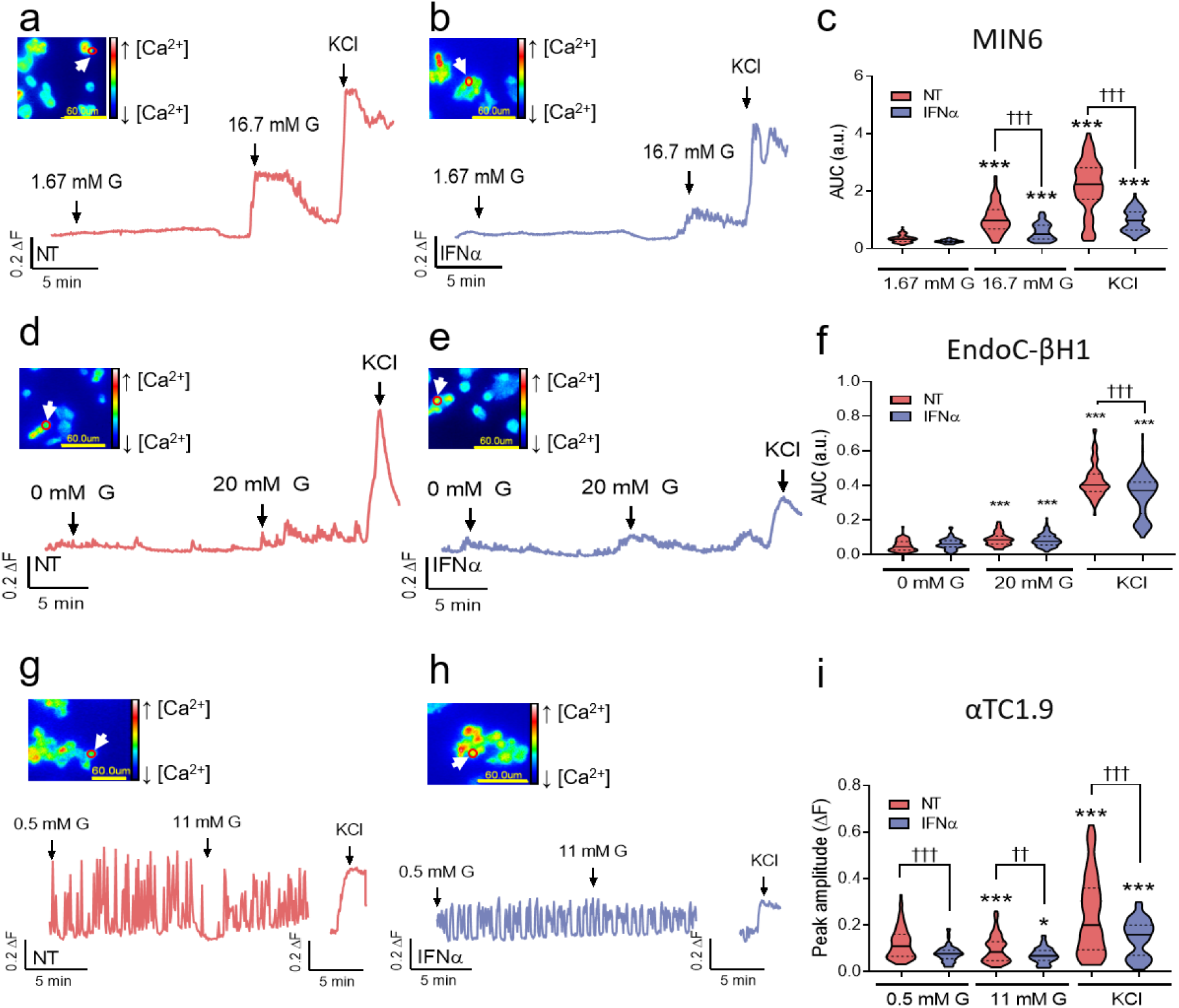
IFNα impairs intracellular Ca^2+^ oscillations in beta and alpha cell models. MIN6 (**a-c**), EndoC-βH1 (**d-f**) and αTC1.9 (**g-i**) cells were left untreated (NT; red) or treated with IFNα (1000 U/ml; blue) for 24 h. Representative recordings of intracellular Ca^2+^ from individual MIN6 (**a, b**), EndoC-βH1 (**d, e**) and αTC1.9 (**g, h**) cells exposed to the glucose concentrations indicated above the traces and to 30 mmol/l KCl. Cells corresponding to representative traces are indicated by a red circle and white arrow in the accompanying image. Scale bar, 60 μm. (**c, f**) Quantification of AUC in intracellular Ca^2+^ changes in MIN6 (**c**) and EndoC-βH1 cells (**f**). (**i**) Ca^2+^ peak amplitude in αTC1.9 cells. Violin plots show the distribution, median (solid lines) and quartiles (dashed lines). (**c**) 82 individual registers (NT: *n*=6 recordings; IFNα: *n*=6 recordings) performed on three separate days. (**f**) 94-109 individual registers (NT: *n*=13 recordings; IFNα: *n*=10 recordings) performed on six separate days. (**i**) 69-80 individual registers (NT: *n*=10 recordings; IFNα: *n*=7 recordings) performed on four separate days. \**p*≤0.05 and \*\*\**p*≤0.001 vs low glucose within the same treatment group; ^††^*p*≤0.01 and ^†††^*p*≤0.001 as indicated by bars; Mixed-effects analysis followed by Tukey’s multiple-comparisons test.

In line with the observations in primary beta cells, IFNα impaired Ca^2+^ oscillations in both beta cell lines. In MIN6 cells, IFNα reduced AUC in response to 16.7 mmol/l glucose and KCl (Fig. 4a-c). KCl-induced Ca^2+^ responses were also attenuated in EndoC-βH1 cells (Fig. 4d-f), demonstrating impaired beta cell Ca^2+^ signalling in both murine and human cell models.

IFNα also disrupted intracellular Ca^2+^ oscillations in αTC1.9 cells (Fig. 4g-i, ESM Fig. 4). In these cells, increasing glucose from 0.5 to 11 mmol/l reduced Ca^2+^ oscillation frequency and peak amplitude in untreated cells (Fig. 4g-i, ESM Fig. 4). Following IFNα exposure, the inhibitory effect of glucose on Ca^2+^ oscillations was attenuated, with nearly 50% reduction in glucose-mediated suppression of oscillation frequency (ESM Fig. 4a). Moreover, IFNα decreased Ca^2+^ peak amplitude under all conditions tested, including stimulation with 0.5 mmol/l glucose, 11 mmol/l glucose and KCl (Fig. 4i). Finally, a single-cell inhibition index confirmed reduced suppression by high glucose following IFNα exposure (ESM Fig. 4b).

Together, these results suggest that IFNα impairs intracellular Ca^2+^ oscillations in alpha and beta cell models, although the functional consequences differ between cell types.

### IFNα reduces SERCA2 abundance and SERCA activity in beta cells but not in alpha cells

We next investigated whether altered SERCA2 abundance or SERCA activity could contribute to impaired Ca^2+^ oscillations (Fig. 5). IFNα reduced SERCA2 protein abundance by approximately 45-50% in both MIN6 and EndoC-βH1 cells, whereas no changes were detected in αTC1.9 cells (Fig. 5a-d, ESM Fig. 5a, b).

**Fig. 5.**
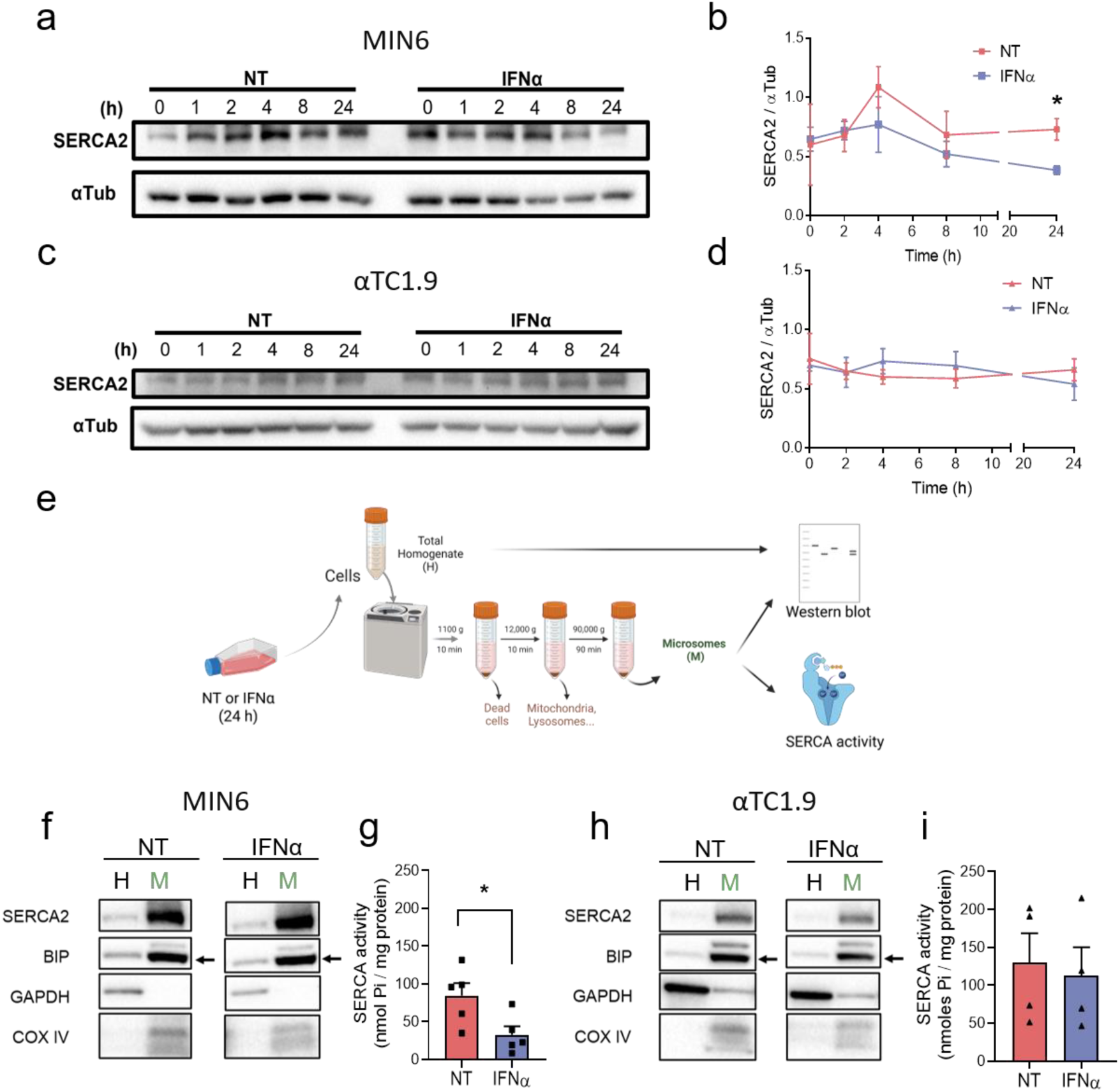
IFNα reduces SERCA2 abundance and SERCA activity in MIN6 but not αTC1.9 cells. MIN6 (**a, b, f, g**; squares) and αTC1.9 (**c, d, h, i**; triangles) cells were left untreated (NT; red) or treated with IFNα (1000 U/ml; blue) for the indicated periods (**a-d**) or for 24 h (**f-i**). (**a, c**) Representative immunoblots of SERCA2 abundance after 0, 1, 2, 4, 8 and 24 h of IFNα exposure, with α-tubulin (αTub) used as loading control. (**b, d**) Quantification of SERCA2 expression in MIN6 (**b**) and αTC1.9 (**d**) cells. Results are means ± SEM of four (**a, b**) or three (**c, d**) independent experiments. \**p*≤0.05 vs NT; paired Student’s *t* test. (**e**) Schematic representation of microsome-enriched fraction isolation. (**f, h**) Immunoblot analysis of SERCA2, BiP, GAPDH and COX IV in whole-cell homogenates (H) and microsome-enriched fractions (M) from MIN6 (**f**) and αTC1.9 (**h**) cells. (**g, i**) SERCA activity in microsome-enriched fractions from MIN6 (**g**) and αTC1.9 (**i**) cells. Results are means ± SEM of 4-5 independent experiments. \**p*≤0.05 vs NT; paired Student’s *t* test.

To determine whether reduced SERCA2 expression was associated with functional impairment, SERCA activity was assessed in microsome-enriched fractions (Fig. 5e), in which SERCA2 enrichment was confirmed by western blotting (Fig. 5f, h). Consistent with the reduced protein expression, SERCA activity was decreased by almost 60% in IFNα-treated MIN6 cells, while no changes were observed in αTC1.9 cells (Fig. 5g, i). In EndoC-βH1 cells, IFNα reduced the amplitude of thapsigargin-induced ER Ca^2+^ release by almost 25%, consistent with impaired SERCA function (ESM Fig. 5c, d).

These findings associate impaired beta cell Ca^2+^ signalling with reduced SERCA2 abundance and SERCA function in MIN6 and EndoC-βH1 but not in αTC1.9 cells.

### IFNα impairs mitochondrial respiration in alpha cells but not in beta cells

As the effects of IFNα on alpha cell Ca^2+^ signalling could not be explained by altered SERCA expression or activity, we investigated whether mitochondrial bioenergetics was affected by IFNα exposure (Fig. 6, ESM Fig. 6).

**Fig. 6.**
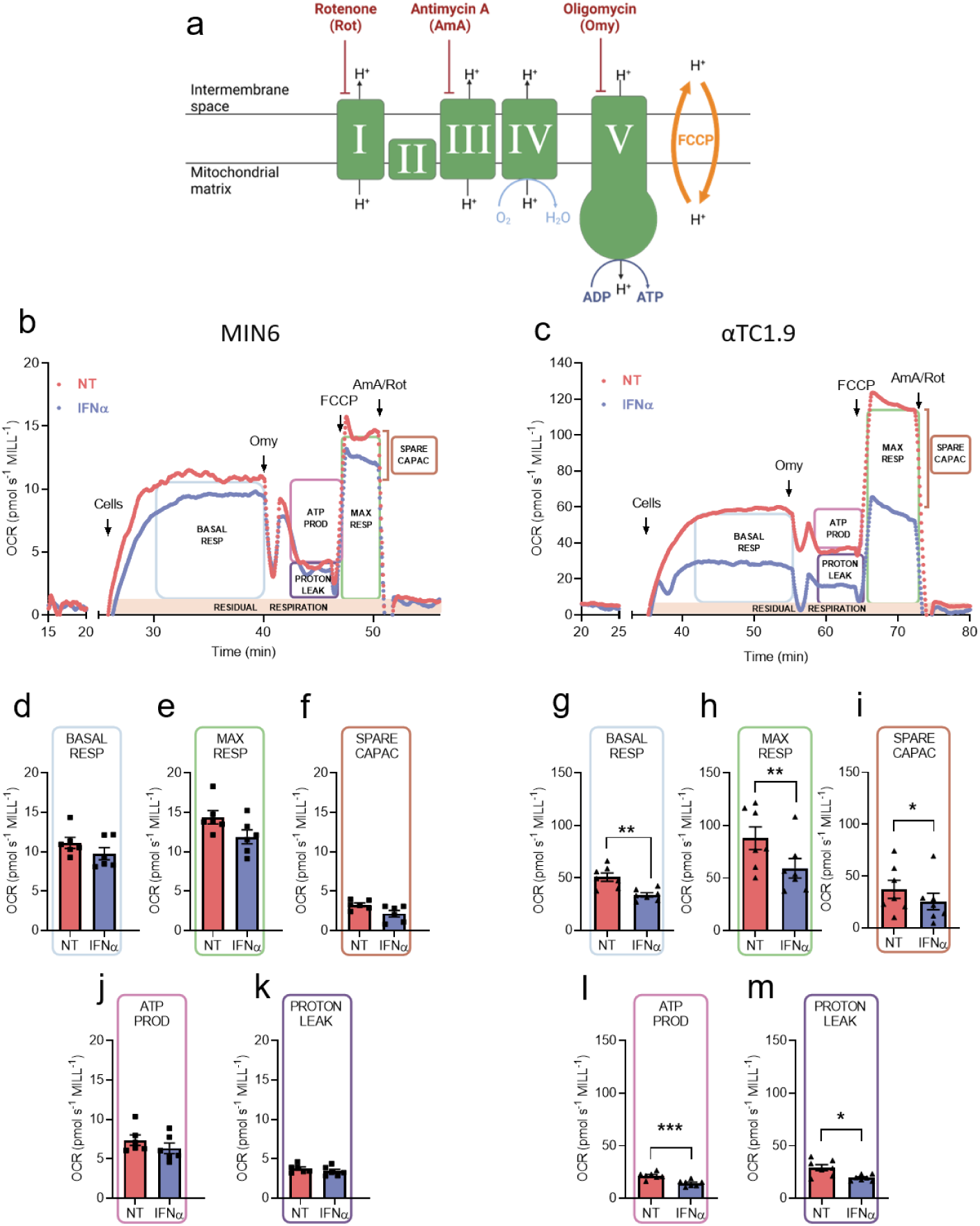
IFNα impairs mitochondrial respiration in αTC1.9 but not MIN6 cells. (**a**) Schematic representation of mitochondrial respiratory-chain complexes I-V in the inner mitochondrial membrane. Inhibition of Complex I by rotenone (Rot), Complex III by antimycin A (AmA) and Complex V by oligomycin (Omy) is shown in red; mitochondrial uncoupling by FCCP is shown in orange. MIN6 (**b, d-f, j, k**; squares) and αTC1.9 (**c, g-i, l, m**; triangles) cells were left untreated (NT; red) or treated with IFNα (1000 U/ml; blue) for 24 h. Representative OCR traces from MIN6 (**b**) and αTC1.9 (**c**) cells. Boxes indicate basal respiration, ATP-linked respiration, proton leak, maximal respiration, spare respiratory capacity and residual oxygen consumption. Arrows indicate additions to the chamber. (**d-m**) Basal respiration (**d, g**), maximal respiration (**e, h**), spare respiratory capacity (**f, i**), ATP-linked respiration (**j, l**) and proton leak (**k, m**). Results are means ± SEM of 6-7 independent experiments. \**p*≤0.05, \*\**p*≤0.01 and \*\*\**p*≤0.001 vs NT; paired Student’s *t* test.

Under untreated conditions, αTC1.9 cells exhibited higher basal and maximal respiration, spare respiratory capacity, ATP-linked respiration and proton leak than MIN6 cells (Fig. 6b-m). Interestingly, IFNα reduced all respiratory parameters by nearly 30% in αTC1.9 cells (Fig. 6g-m) but had no significant effects in MIN6 (Fig. 6a-f) or EndoC-βH1 cells (ESM Fig. 6).

### Mitochondrial complex expression differs between αTC1.9 and MIN6 cells but is not altered by IFNα

We then examined whether differences in respiratory capacity or the effects of IFNα were associated with mitochondrial respiratory-chain complex abundance (Fig. 7, ESM Fig. 7).

**Fig. 7.**
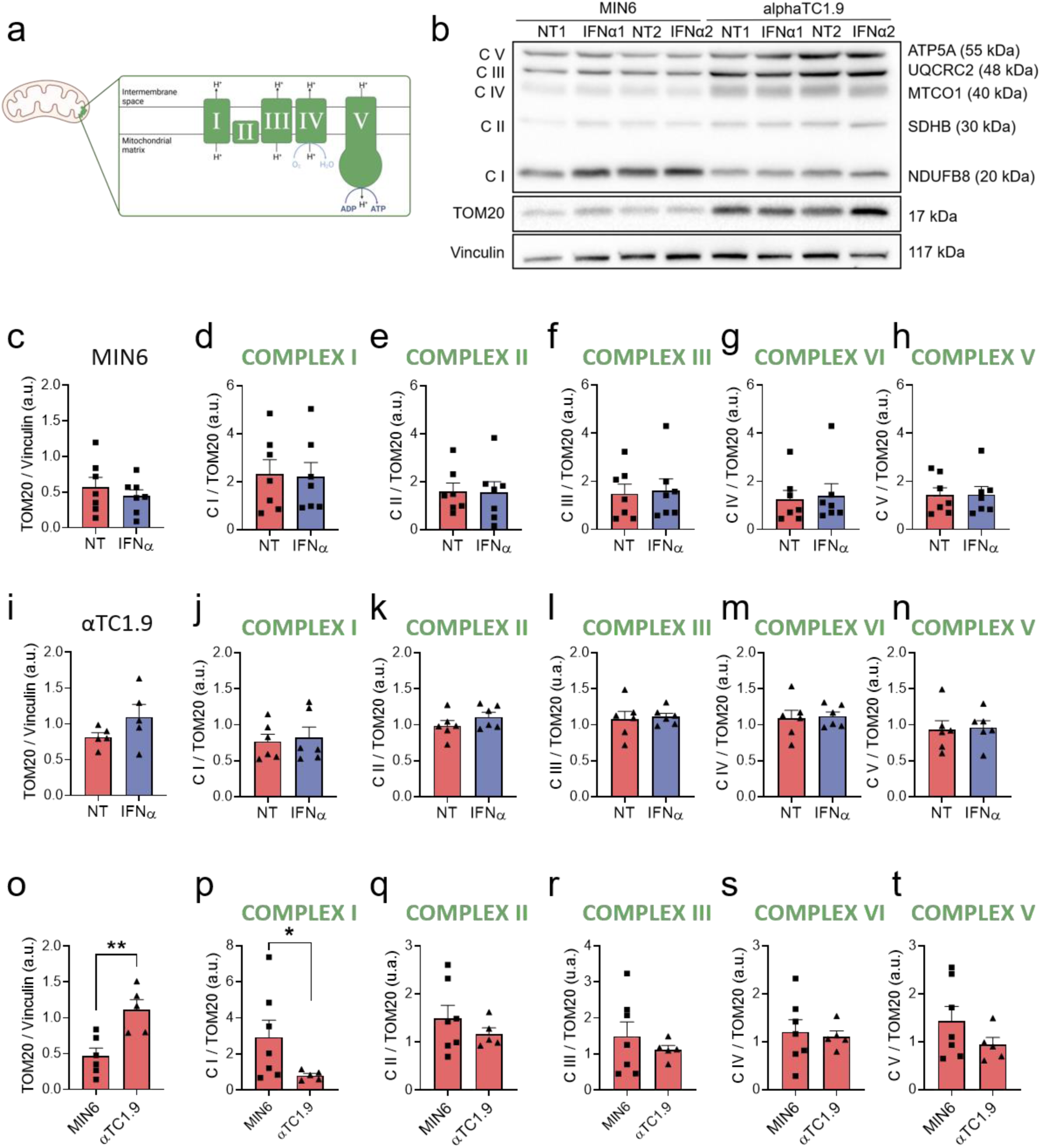
Mitochondrial respiratory-chain complex abundance differs between MIN6 and αTC1.9 cells but is not altered by IFNα. (**a**) Schematic representation of mitochondrial respiratory-chain complexes I-V in the inner mitochondrial membrane. MIN6 (**b-h, o-t**; squares) and αTC1.9 (**b, i-t**; triangles) cells were left untreated (NT; red) or treated with IFNα (1000 U/ml; blue) for 24 h. (**b**) Representative immunoblots of respiratory-chain Complexes I-V, TOM20 and vinculin in MIN6 and αTC1.9 cells. (**c-n**) Quantification of TOM20 and Complexes I-V in MIN6 (**c-h**) and αTC1.9 (**i-n**), normalised to vinculin. (**o-t**) Comparison of TOM20 abundance (**o**) and Complexes I-V (**p-t**) between the two cell lines, with respiratory-complex abundance normalised to TOM20. Results are means ± SEM of 5-7 independent experiments. \**p*≤0.05 and \*\**p*≤0.01 as indicated by bars; unpaired Student’s *t* test.

αTC1.9 cells expressed higher levels of mitochondrial respiratory chain complexes than MIN6 cells when protein abundance was normalised to vinculin, except for Complex I (ESM Fig. 7k-o). This observation led us to examine whether differences in mitochondrial mass contributed to the distinct respiratory profiles of the two cell types.

Mitochondrial mass was assessed by measuring the outer mitochondrial membrane protein TOM20. αTC1.9 cells had 2.4-fold greater TOM20 abundance than MIN6 cells (Fig. 7c, i), consistent with greater mitochondrial mass and respiratory activity. After normalisation to TOM20, most differences in respiratory-complex abundance disappeared (Fig. 7o-t). However, Complex I expression remained significantly higher in MIN6 cells, exhibiting a 3.6-fold greater abundance per unit mitochondrial mass than αTC1.9 cells (Fig. 7p).

Notably, IFNα did not change TOM20 or respiratory-complex protein levels when normalised to vinculin or TOM20 (Fig. 7, ESM Fig. 7).

These results indicate greater mitochondrial mass in αTC1.9 cells and relative Complex I enrichment in MIN6 cells but exclude changes in mitochondrial mass or complex abundance as explanations for the IFNα-induced respiratory phenotype.

### IFNα selectively impairs NADH-linked respiration in alpha cells

To investigate whether the effects of IFNα on mitochondrial respiration were linked to alterations in specific electron transfer pathways, NADH- and succinate-linked respiration were assessed separately in digitonin-permeabilised MIN6 and αTC1.9 cells (Fig. 8).

**Fig. 8.**
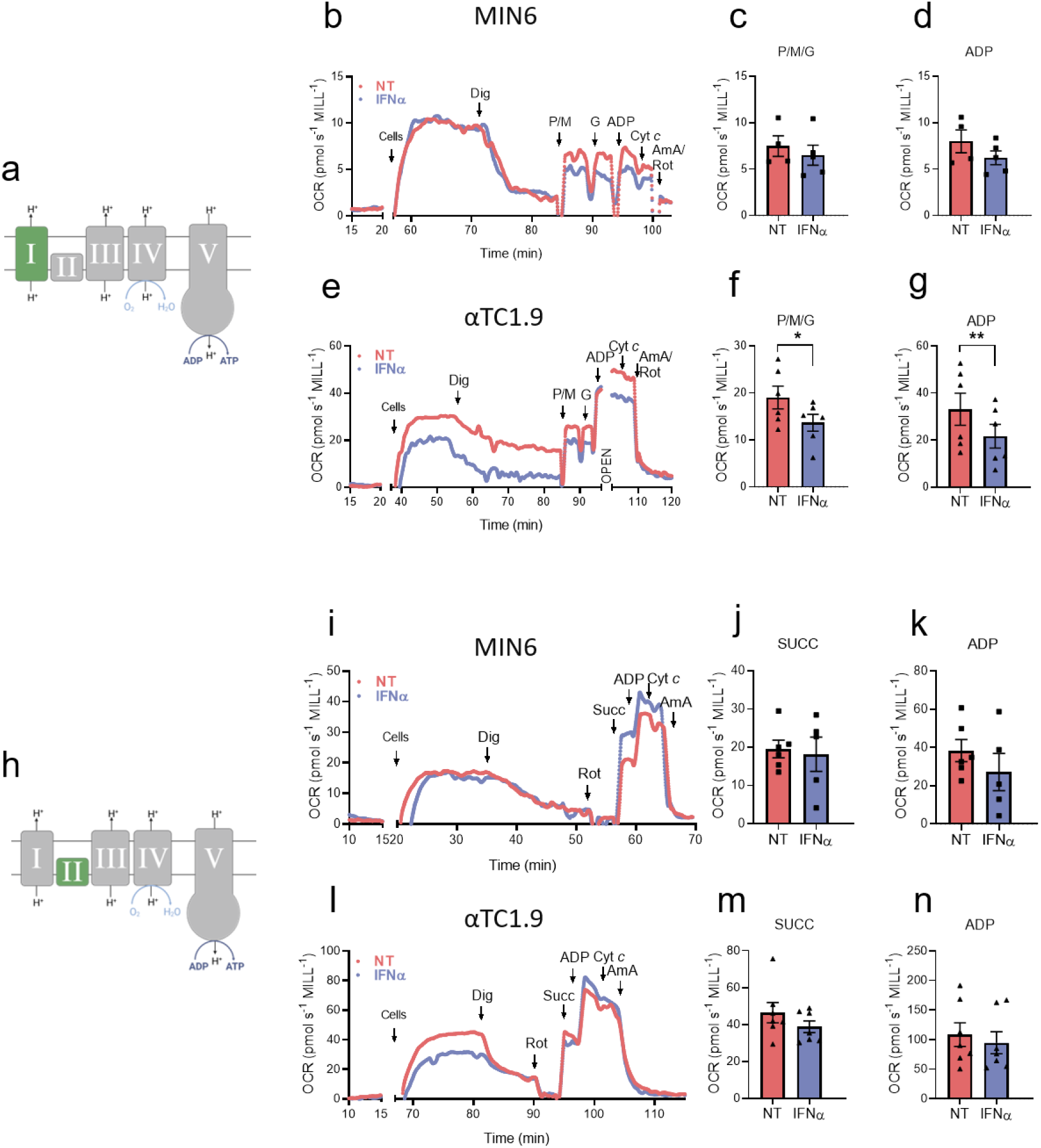
IFNα selectively impairs NADH-linked respiration in αTC1.9 cells. (**a, h**) Schematics of mitochondrial respiratory-chain complexes I-V, highlighting the Complex I (**a**) and Complex II (**h**). MIN6 (**b-d, i-k**; squares) and αTC1.9 (**e-g, l-n**; triangles) cells were left untreated (NT; red) or treated with IFNα (1000 U/ml; blue) for 24 h. Representative OCR traces from MIN6 (**b, i**) and αTC1.9 (**e, l**) cells during assessment of NADH-linked (**b, e**) and succinate-linked (**i, l**) respiration. Arrows indicate additions to the chamber. NADH-linked respiration was quantified following addition of pyruvate (P), malate (M) and glutamate (G) (**c, f**) and ADP (**d, g**). Succinate-linked respiration was quantified following addition of succinate (**j, m**) and ADP (**k, n**) in the presence of rotenone (Rot). Results are means ± SEM of 4-7 independent experiments. \**p*≤0.05 and \*\**p*≤0.01 vs NT; unpaired Student’s *t* test. Dig: Digitonin; Cyt *c*: Cytochrome *c*.

In MIN6 cells, IFNα did not affect respiration supported by Complex I substrates under either substrate-supported or ADP-stimulated conditions (Fig. 8b-d). Similarly, IFNα did not alter Complex II-supported respiration following the addition of succinate in the presence of rotenone (Fig. 8i-k). In contrast, in αTC1.9 cells, IFNα reduced pyruvate-, malate-, and glutamate-supported respiration by approximately 28% and ADP-stimulated oxidative phosphorylation by 13% (Fig. 8e-g). Succinate-linked respiration, however, remained unchanged (Fig. 8l-n).

These findings indicate that IFNα selectively impairs NADH-linked mitochondrial respiration in alpha cells without affecting succinate-linked respiration.

## Discussion

In the present study, we show that acute IFNα exposure disrupts glucose-regulated hormone secretion in vivo, ex vivo and in endocrine cell models. IFNα impaired GSIS and attenuated the regulation of glucagon secretion by glucose and insulin. These changes were accompanied by defective intracellular Ca^2+^ signalling in beta and alpha cells. However, the associated mechanisms differed between cell types: beta cell dysfunction was associated with reduced SERCA2 abundance and SERCA activity, whereas alpha cell dysfunction was accompanied by impaired mitochondrial respiration, predominantly affecting the NADH-linked pathways. These effects occurred without detectable changes in viability, ROS production, proliferation or endocrine cell composition, indicating functional impairment rather than a cytotoxic phenotype.

Previous studies have established IFNα as a key component of early islet inflammation in type 1 diabetes [2, 7, 8, 10]. Our findings extend these established immunological effects by showing that IFNα also interferes with stimulus-secretion coupling. Thus, alterations in endocrine function may represent an additional and previously underappreciated component of IFNα-mediated islet dysfunction.

Acute systemic IFNα administration altered glucose and hormonal responses during the IPGTT. IFNα increased glucose AUC overall, although sex-stratified comparisons detected a significant increase only in male mice. In females, IFNα did not significantly alter glucose AUC. However, untreated females already exhibited a pronounced glucose excursion at 15 min, and both groups returned towards basal glucose concentrations by 120 min. IFNα-treated females also showed lower basal glucagon concentrations, while serum insulin concentrations at baseline and 120 min were not significantly altered in either sex. Although the change in circulating insulin between baseline and 120 min was reduced following IFNα treatment, the absence of early post-glucose measurements prevents us from drawing conclusions about the acute insulin secretory response. These findings indicate that acute IFNα exposure perturbs glucose regulation in vivo, although the available hormone measurements do not identify the endocrine or extra-pancreatic mechanisms underlying this phenotype. Consistent with a broader systemic effect, Koivisto et al. also reported impaired glucose tolerance following acute IFNα administration in healthy individuals, accompanied by increased circulating insulin, glucagon and other counterregulatory hormones [35]. The endocrine responses observed in our study differed from those reported in humans, in whom glucose intolerance occurred despite increased circulating insulin. This difference may reflect species-specific responses as well as differences in experimental design. Altered insulin sensitivity, hepatic glucose metabolism, counterregulatory hormone action or insulin clearance may have contributed to the in vivo phenotype. The defects observed in isolated islets support a direct islet contribution, even though the in vivo phenotype cannot be attributed exclusively to endocrine cell dysfunction.

Despite impaired beta cell Ca^2+^ signals, the effects of IFNα on insulin secretion were modest and depended on the experimental model. Insulin secretion was preserved in MIN6 cells, whereas isolated islets showed reduced responses to high glucose and IBMX-potentiated stimulation. This pattern suggests that acute IFNα exposure may impair beta cell functional reserve, with secretory defects emerging mainly under increased secretory demand. Preservation of insulin secretion in MIN6 cells is consistent with previous reports showing that IFNα does not impair GSIS in human beta cell models, despite inducing inflammatory and stress responses [36–38].

The dissociation between impaired Ca^2+^ oscillations and preserved secretion in MIN6 cells may reflect compensatory mechanisms downstream of Ca^2+^ influx. Reduced SERCA2 abundance and SERCA activity may contribute to the altered Ca^2+^ dynamics, given the role of SERCA2 in ER Ca^2+^ homeostasis and beta cell Ca^2+^ responses [39]. The reduction in SERCA2 levels in EndoC-βH1 cells was similarly accompanied by a smaller thapsigargin-induced Ca^2+^ response, consistent with impaired ER Ca^2+^ handling in a human beta cell model. This is in line with evidence that cytokine exposure and ER stressors can disrupt intracellular Ca^2+^ dynamics without proportionally affecting insulin secretion [40]. However, the association between reduced SERCA2 and altered Ca^2+^ dynamics remains correlative, as we did not test whether restoring SERCA2 expression or activity rescues the Ca^2+^ phenotype. Compensation downstream of Ca^2+^, potentially involving the metabolic amplifying pathway, may help preserve insulin secretion in MIN6 cells, but was not directly assessed.

IFNα also directly disrupted alpha cell function. Rather than causing uniform hypersecretion or hyposecretion, IFNα narrowed the glucose-responsive range of glucagon secretion. Secretion was reduced under stimulatory low-glucose conditions, while suppression by higher glucose concentrations was attenuated. In αTC1.9 cells, IFNα also reduced insulin-mediated inhibition without altering glucagon content. This phenotype resembles aspects of the dysfunction reported in islets from non-diabetic GADA-positive donors, which retain insulin secretory responsiveness but show impaired glucose-mediated suppression of glucagon secretion and alpha cell alterations in metabolic pathways [17]. Compromised α-cell function has also been documented in established type 1 diabetes [16], with more recent integrated electrophysiological and transcriptomic analyses identifying increased excitability and exocytosis, loss of glucose-mediated suppression, and extensive metabolic reprogramming in alpha cells from type 1 diabetes donors, further supporting a link between immune and metabolic alterations and alpha cell dysfunction [19]. Our findings therefore identify IFNα as a potential contributor to alpha cell dysfunction during early islet autoimmunity.

In αTC1.9 cells, the secretory and Ca^2+^ abnormalities were accompanied by impaired mitochondrial respiration. This was not associated with changes in TOM20 or respiratory-chain complex abundance, supporting a functional defect rather than loss of mitochondrial mass or respiratory proteins. Previous transcriptomic analyses of IFNα-exposed human induced pluripotent stem cell-derived islet-like cells identified depletion of gene sets related to oxidative phosphorylation and respiratory electron transport [15]. Moreover, alpha cells from donors with type 1 diabetes exhibited alterations in carbohydrate metabolism and the TCA cycle, and transcripts related to mitochondrial respiration were associated with their abnormal electrophysiological phenotype [19]. Our results extend these transcriptomic findings by demonstrating reduced oxygen consumption, predominantly affecting NADH-linked (Complex I-supported) respiration in αTC1.9 cells. Importantly, reduced respiration supported by pyruvate, malate and glutamate does not demonstrate direct inhibition of Complex I, as altered substrate transport, matrix dehydrogenase activity or NADH generation could produce the same phenotype.

In conclusion, acute IFNα exposure disrupts endocrine cell stimulus-secretion coupling, associated with SERCA2-related Ca^2+^ dysregulation in beta cells and impaired NADH-linked mitochondrial respiration in alpha cells. These findings extend the pathogenic role of IFNα beyond immune activation and beta cell immunogenicity to dysregulation of insulin and glucagon secretion.

## Acknowledgements

The authors are grateful to Beatriz Bonmati Botella, Maria Luisa Navarro, and Salomé Ramon from the Universidad Miguel Hernández de Elche, Spain, for their excellent technical support. The authors would like to thank Dr. Pablo M. Garcia-Roves from the University of Barcelona for his insightful discussions regarding the mitochondrial respiration results.

During the preparation of this work, the authors used Microsoft Copilot to improve the language and readability of the manuscript. After using this tool, the authors reviewed and edited the content as needed, and they take full responsibility for the content of the publication.

## Author contributions

LM and RSdS conceived and supervised the study; AAP-S, DG-L, RSdS and LM set up methods and models, performed the experiments, acquired, analysed and interpreted the data, and wrote the original draft of the article; FMG and FR provided resources; all the authors reviewed, edited and approved the final version of the manuscript; LM acquired funding. LM is the guarantor of this work and, as such, has full access to all the data in the study and take responsibility for the integrity of the data and the accuracy of the data analysis.

