## Supplementary Material for "IFNα differentially disrupts stimulus-secretion coupling in pancreatic alpha and beta cells"

**Electronic Supplementary Material**

### ESM Methods

#### Animals

C57BL/6 and Glu-Venus mice were housed under standard conditions with a 12 h light-dark cycle and ad libitum access to food and water. For ex vivo experiments, mice were euthanised by cervical dislocation before pancreatic islet isolation. Mice undergoing IPGTT were euthanised by $CO_2$ overdose before islet isolation.

#### Islet isolation and dispersion

Pancreatic islets were isolated from male and female mice by collagenase digestion following pancreatic duct perfusion. Pancreases were digested with collagenase (Sigma-Aldrich) at 37°C and diluted in ice-cold HEPES-buffered Krebs-Ringer solution containing (mmol/l): 115 NaCl, 5 KCl, 10 $NaHCO_3$, 1.1 $MgCl_2$, 1.2 $NaH_2PO_4$, 2.5 $CaCl_2$, 25 HEPES, and 5 glucose, supplemented with 0.25% BSA (pH 7.4). Islets were hand-picked under a stereomicroscope and allowed to recover for 2 h at 37°C before experimental treatment or dispersion.

For primary cell cultures, islets were dispersed with 0.25% trypsin-EDTA for 3 min at 37°C. Cells were washed, centrifuged and seeded onto poly-L-lysine-coated plates in RPMI 1640 medium supplemented with 8% heat-inactivated FBS, 2 mmol/l L-glutamine, 100 U/ml penicillin and 100 µg/ml streptomycin. Cells were maintained at 37°C in a humidified atmosphere containing 5% $CO_2$ and allowed to attach for 48 h before experimental treatments.

#### Cell culture

MIN6 cells (RRID: CVCL_0431) were cultured in high-glucose DMEM supplemented with 15% FBS, 0.05 mmol/l 2-mercaptoethanol, 100 U/ml penicillin and 100 µg/ml streptomycin [1].

αTC1.9 cells (RRID: CVCL_0150) were maintained in phenol red-free DMEM supplemented with 10% FBS, 0.1 mmol/l non-essential amino acids, 15 mmol/l HEPES, 2 mmol/l L-glutamine, 19 mmol/l $NaHCO_3$, 100 U/ml penicillin and 100 µg/ml streptomycin, with glucose adjusted to a final concentration of 16 mmol/l [2].

EndoC-βH1 cells (RRID: CVCL_L909) were cultured in low-glucose DMEM supplemented with 5.5 µg/ml transferrin, 6.7 ng/ml sodium selenite, 10 mmol/l nicotinamide, 0.05 mmol/l 2-mercaptoethanol, 2% fatty acid-free BSA, 100 U/ml penicillin and 100 µg/ml streptomycin.

Cells were seeded on Matrigel/fibronectin-coated culture plates as previously described [3, 4]. For experimental treatments, the culture medium was supplemented with 2% FBS.
All cell lines were cultured at 37°C in a humidified atmosphere containing 5% $CO_2$. Moreover, they were routinely tested and confirmed negative for *Mycoplasma* contamination using the MycoAlert Mycoplasma Detection Kit (Lonza, Basel, Switzerland).

**DCF assay**
Oxidative stress was assessed using the fluorescent probe 2′,7′-dichlorofluorescein diacetate (DCF, ref: 278475, Sigma-Aldrich, St. Louis, USA) as previously described [5]. Briefly, cells were seeded in 96-well black plates and loaded with 10 μM DCF for 30 min at 37 °C, then washed with PBS. DCF fluorescence was quantified in a POLASTAR plate reader (BMG Labtech, Germany). Data is expressed as DCF fluorescence normalized by total protein, measured by Bradford assay.

**Immunofluorescence**
Immunofluorescence was performed as described [6, 7]. Briefly, cells washed with cold PBS were fixed with 4% paraformaldehyde, permeabilized, and incubated overnight with primary antibodies against insulin and KI67. The Alexa Fluor 647 polyclonal goat anti-mouse IgG and the Alexa Fluor 546 polyclonal goat anti-rabbit IgG were used as secondary antibodies for insulin and KI67 staining, respectively. Glucagon-positive cells were identified by green fluorescence. After nuclei staining with Hoechst 33342, coverslips were mounted using a fluorescent mounting medium (Dako, Carpintera, CA, USA), and images were acquired on an IN-Cell Analyzer 6000 Cell Imaging system (GE Healthcare, Little Chalfont, UK) with a x20 objective. At least 25 images per coverslip were analysed, and at least 2,000 total cells/animal/condition were counted. Quantification was performed using the open-source FIJI (version 2.0; https://fiji.sc) software.

**Cell viability assessment**
The percentage of apoptosis was assessed after incubation with the DNA-binding dyes Hoechst 33342 (5 μg/ml; Sigma-Aldrich) and propidium iodide (10 μg/ml; Sigma-Aldrich) using fluorescence microscopy [8, 9]. At least 600 cells were counted per experimental condition by two independent researchers, one of whom was blinded to sample identity, with >90% agreement between observers. Apoptosis was expressed as the percentage of apoptotic cells relative to the total number of cells.

**Intracellular $Ca^{2+}$ recordings**

Intracellular $Ca^{2+}$ dynamics were measured in intact islets or cultured cells plated on glass coverslips. EndoC-βH1, MIN6 and αTC1.9 cells and intact islets were loaded with 2 μmol/l Fura-2/AM (Sigma-Aldrich) for 1 h at room temperature in a humidified chamber protected from light, as previously described [7, 10]. Fluorescence was recorded every 2 s using an Axiovert 200M inverted microscope (Zeiss, Jena, Germany) equipped with a TILL Photonics polychromator for alternating excitation at 340 and 380 nm using 10-nm bandpass filters (Omega Optical, Brattleboro, VT, USA), and a Hamamatsu EMC9100 camera (Hamamatsu Photonics, Hamamatsu, Japan). Images were analysed using Aquacosmos software (version 2.6; Hamamatsu Photonics).

Primary alpha cell $Ca^{2+}$ dynamics were assessed in intact islets from C57BL/6 mice loaded with 2 μmol/l Fluo-4/AM (Invitrogen, Waltham, MA, USA) under the same loading conditions [11, 12]. Optical sections of 4.9 μm were acquired every 2 s using an Axiovert 200M microscope coupled to a Zeiss LSM 510 confocal module with an 8 μm pinhole and a 40x objective. Fluo-4 was excited using a 488 nm argon laser, and fluorescence emission was collected using a 505-530 nm band-pass filter and analysed using ZEN 2009 software (Zeiss). Primary alpha cells were identified according to previously established criteria [11, 13, 14]: (1) localisation within the islet mantle; (2) a characteristic high-frequency $Ca^{2+}$ oscillatory pattern at low glucose concentrations that was suppressed by high glucose; and (3) a positive $Ca^{2+}$ response to adrenaline.

Preparations were perfused at 2 ml/min with Krebs-Ringer bicarbonate buffer containing (mmol/l): 141 NaCl, 5.5 KCl, 1 $MgCl_2$, 2 $CaCl_2$ and 20 HEPES (pH 7.4).

Basal fluorescence was recorded for 5 min at 0 mmol/l glucose for EndoC-βH1 cells, 1.67 mmol/l for MIN6 cells, 3 mmol/l for intact islets, and 5.6 mmol/l for primary alpha cells and αTC1.9 cells. Preparations were subsequently exposed to 20 mmol/l glucose for EndoC-βH1 cells, 16.7 mmol/l for MIN6 cells, 8 or 11 mmol/l for intact islets, and 0.5 mmol/l followed by 11 mmol/l for primary alpha cells and αTC1.9 cells. Each condition was recorded for 10 min.

Glucose-independent $Ca^{2+}$ responses were elicited with 30 mmol/l KCl in EndoC-βH1, MIN6 and αTC1.9 cells and intact islets, or with 5 μmol/l adrenaline in primary alpha cells. Fura-2 responses were expressed as the 340/380 fluorescence ratio, whereas Fluo-4 responses were expressed as $F/F_0$.

In EndoC-βH1 and MIN6 cells and intact islets, $Ca^{2+}$ responses were quantified as the area under the curve during the last 5 min of stimulation relative to basal fluorescence and expressed in arbitrary units. Peak amplitude was additionally determined in intact islets exposed to 8 mmol/l glucose and in αTC1.9 cells under all stimulation conditions. Oscillation frequency was quantified in primary alpha and αTC1.9 cells and expressed as peaks/min.

For each αTC1.9 cell recording, the glucose inhibition index was calculated as: 1 - (mean peak amplitude at 11 mmol/l glucose / mean peak amplitude at 0.5 mmol/l glucose), using the final 5 min of each stimulation period.

The KCl stimulation index was calculated as: mean peak amplitude during KCl stimulation / mean peak amplitude during 0.5 mmol/l glucose, using paired measurements from each cell.

Data were analysed using GraphPad Prism 8 (GraphPad Software, San Diego, CA, USA).

**Hormone secretion and content**

Insulin and glucagon secretion were assessed using a static incubation assay as previously described [1, 2, 15]. For intact islets, size-matched islets were hand-picked and distributed into groups of ten.

After 24 h of treatment, cells and islets were washed with Krebs-Ringer bicarbonate (KRB) buffer containing (mmol/l): 120 NaCl, 5.4 KCl, 1.2 $KH_2PO_4$, 1.2 $MgSO_4$, 2.4 $CaCl_2$, 20 HEPES (pH 7.4), supplemented with 0.1 % BSA. MIN6 and αTC1.9 cells were preincubated for 2 h in KRB containing 0 or 5.6 mmol/l glucose, respectively. Islets were preincubated for 1 h in KRB containing 1 mmol/l glucose.

Cells were subsequently incubated for 30 min and islets for 1 h under the following conditions: MIN6 cells, 1.67 mmol/l glucose, 16.7 mmol/l glucose, or 16.7 mmol/l glucose plus 30 mmol/l KCl; αTC1.9 cells, 0.5 mmol/l glucose, 11 mmol/l glucose, or 0.5 mmol/l glucose plus 10 nmol/l insulin; and islets, 1 mmol/l glucose, 5.6 mmol/l glucose, 16.7 mmol/l glucose, or 16.7 mmol/l glucose plus 50 µmol/l 3-isobutyl-1-methylxanthine (IBMX).

Supernatants were collected on ice, and insulin or glucagon concentrations were measured by ELISA. Aprotinin was added to glucagon-containing samples at a final concentration of 7 U/ml. For hormone content measurements, cells were washed with ice-cold PBS and extracted with 150 µl ice-cold HCl-ethanol. Islets were transferred to sealed tubes containing the same extraction solution. Samples were incubated for 24 h at 4°C under continuous agitation, after which the supernatants were collected for ELISA analysis. Hormone secretion and intracellular content were measured using mouse or human ELISA kits (Mercodia, Uppsala, Sweden), following the manufacturer’s instructions.

**Subcellular fractionation**

MIN6 and αTC1.9 cells were cultured in 100 mm dishes at densities of 8 x $10^6$ and 5 x $10^6$ cells per dish, respectively. After 24 h of treatment, cells were washed once with ice-cold PBS and scraped into centrifugation buffer containing (mmol/l): 250 sucrose, 2 EGTA, 10 KCl, 1.5 $MgCl_2$, 10 Tris-HCl (pH 7.4), supplemented with 0.1% fatty acid-free BSA and protease inhibitor cocktail. Cell suspensions were homogenised by five passages through a 30-gauge needle, avoiding foam formation throughout the procedure.

An aliquot of the whole-cell homogenate was retained for analysis of SERCA2 abundance by western blotting. The remaining homogenate was sequentially centrifuged at 1,100 *g* for 10 min and 12,000 *g* for 10 min at 4°C, with transfer of the supernatant to a fresh tube after each step. The final supernatant was ultracentrifuged at 90,000 *g* for 90 min at 4°C. The microsome-enriched pellet was resuspended in 50 µl centrifugation buffer and stored at -80°C until analysis.

**SERCA activity**

SERCA activity was assessed in microsome-enriched fractions obtained by subcellular fractionation using a colorimetric ATPase/GTPase Activity Assay Kit (Sigma-Aldrich) according to the manufacturer's instructions. Microsome-enriched fractions containing 10 µg total protein were incubated for 30 min at 37°C with 4 mmol/l ATP and 1.5 mmol/l $CaCl_2$, in the absence or presence of 20 µmol/l cyclopiazonic acid. SERCA activity was calculated as the difference between total ATPase activity and cyclopiazonic acid-insensitive ATPase activity. Liberated inorganic phosphate was quantified colorimetrically by measuring absorbance at 620 nm using an iMark Microplate Reader (Bio-Rad). SERCA activity was expressed as nmol phosphate released per milligram of protein.

In EndoC-βH1 cells, ER $Ca^{2+}$ stores were assessed indirectly from the cytosolic $Ca^{2+}$ response to acute SERCA inhibition, as previously described [16]. Fura-2AM-loaded cells were stimulated with 1 µmol/l thapsigargin, and the resulting $Ca^{2+}$ response was quantified as AUC.

**High-resolution respirometry**

Mitochondrial respiration was assessed in intact and digitonin-permeabilised cells by measuring oxygen consumption rate (OCR) using high-resolution respirometry (Oxygraph-2k; OROBOROS Instruments, Innsbruck, Austria). Cells were detached with 0.25% trypsin-EDTA for 3 min at 37°C and resuspended in MiR05 respiration medium containing (mmol/l): 110 D-sucrose, 0.5 EGTA, 3 $MgCl_2$, 60 lactobionic acid, 20 taurine, 10 $KH_2PO_4$, and 20 HEPES (pH 7.1), supplemented with 0.1% fatty acid-free BSA. Cells in suspension were then transferred to Oxygraph-2k chambers at a density of 3.0 x $10^6$ MIN6 cells/ml, 1.5 x $10^6$ αTC1.9 cells/ml, and 2.0 x $10^6$ EndoC-βH1 cells/ml.

Intact-cell respiration was assessed in all three cell lines, whereas permeabilised-cell experiments were performed only in MIN6 and αTC1.9 cells.

In intact cells, basal respiration was recorded before sequential addition of oligomycin (50 nmol/l), FCCP (0.5 µmol/l for MIN6 and αTC1.9 cells and 1.5 µmol/l for EndoC-βH1 cells), and rotenone (0.5 µmol/l) plus antimycin A (2.5 µmol/l). ATP-linked respiration was calculated as basal respiration minus oligomycin-insensitive respiration. Proton leak was calculated as oligomycin-insensitive respiration minus residual oxygen consumption after rotenone and antimycin A addition. Spare respiratory capacity was calculated as maximal FCCP-stimulated respiration minus basal respiration.

For permeabilised-cell experiments, plasma membranes were permeabilised with digitonin at 6 µg/ml for MIN6 cells and 4 µg/ml for αTC1.9 cells. NADH-linked respiration was assessed by sequential addition of pyruvate (5 mmol/l), malate (2 mmol/l) and glutamate (10 mmol/l), followed by ADP (5 mmol/l) to stimulate oxidative phosphorylation. Succinate-linked respiration was assessed after Complex I inhibition with rotenone (0.5 µmol/l), followed by addition of succinate (10 mmol/l) and ADP (5 mmol/l). Outer mitochondrial membrane integrity was evaluated by addition of cytochrome *c* (10 µmol/l); no substantial increase in OCR was observed following its addition. Experiments were terminated with antimycin A (2.5 µmol/l) alone or antimycin A plus rotenone, according to the protocol.

OCR calculations and data analysis were performed using O2k-Software DatLab 7.4 (OROBOROS Instruments). Control and IFNα-treated samples were alternated between chambers to minimise chamber-specific variability.

**ESM Tables and Figures**

| Antibody | Manufacturer | Catalogue number | Species raised in | Dilution | RRID |
|---|---|---|---|---|---|
| BIP | Cell Signaling | 3177 | Mouse | 1:1000 | AB_2119845 |
| COX IV | Cell Signaling | 4850 | Rabbit | 1:1000 | AB_2085424 |
| GAPDH | Cell Signaling | 5174 | Rabbit | 1:1000 | AB_10622025 |
| OxPhos Rodent WB Antibody cocktail | Invitrogen | 45-8099 | Mouse | 1:1000 | AB_2533835 |
| SERCA2 | Santa Cruz | sc-376235 | Mouse | 1:1000 | AB_10989947 |
| TOM20 | Proteintech | 11802-1-AP | Rabbit | 1:1000 | AB_2207530 |
| Vinculin | Sigma-Aldrich | SAB4200080 | Mouse | 1:1000 | AB_10604160 |
| α-Tubulin | Sigma-Aldrich | T9026 | Mouse | 1:5000 | AB_477593 |
| Goat anti-mouse IgG | Bio-rad | 170-6516 | Goat | 1:5000 | AB_11125547 |
| Goat anti-rabbit IgG | Bio-rad | 170-6515 | Goat | 1:5000 | AB_11125142 |
| Alexa Fluor 568 goat anti-mouse IgG | Invitrogen | A-11031 | Goat | 1:500 | AB_144696 |

**ESM Table 1. Antibody used in the present study.**

| Gene | Forward | Reverse |
|---|---|---|
| | Sequence (5’-3’) | Sequence (5’-3’) |
| ***Hprt1*** | GGTTAAGCAGTACAGCCCCA | TCCAACACTTCGAGAGGTCC |
| ***Cxcl10*** | TCAGGCTCGTCAGTTCTAAGT | CCTTGGGAAGATGGTGGTTAAG |
| ***Mx1*** | CTCACCTCCCACATCTGTAAATC | GTATGTCTGCACCGTACTTCTG |

**ESM Table 2. Primers used in the present study.**

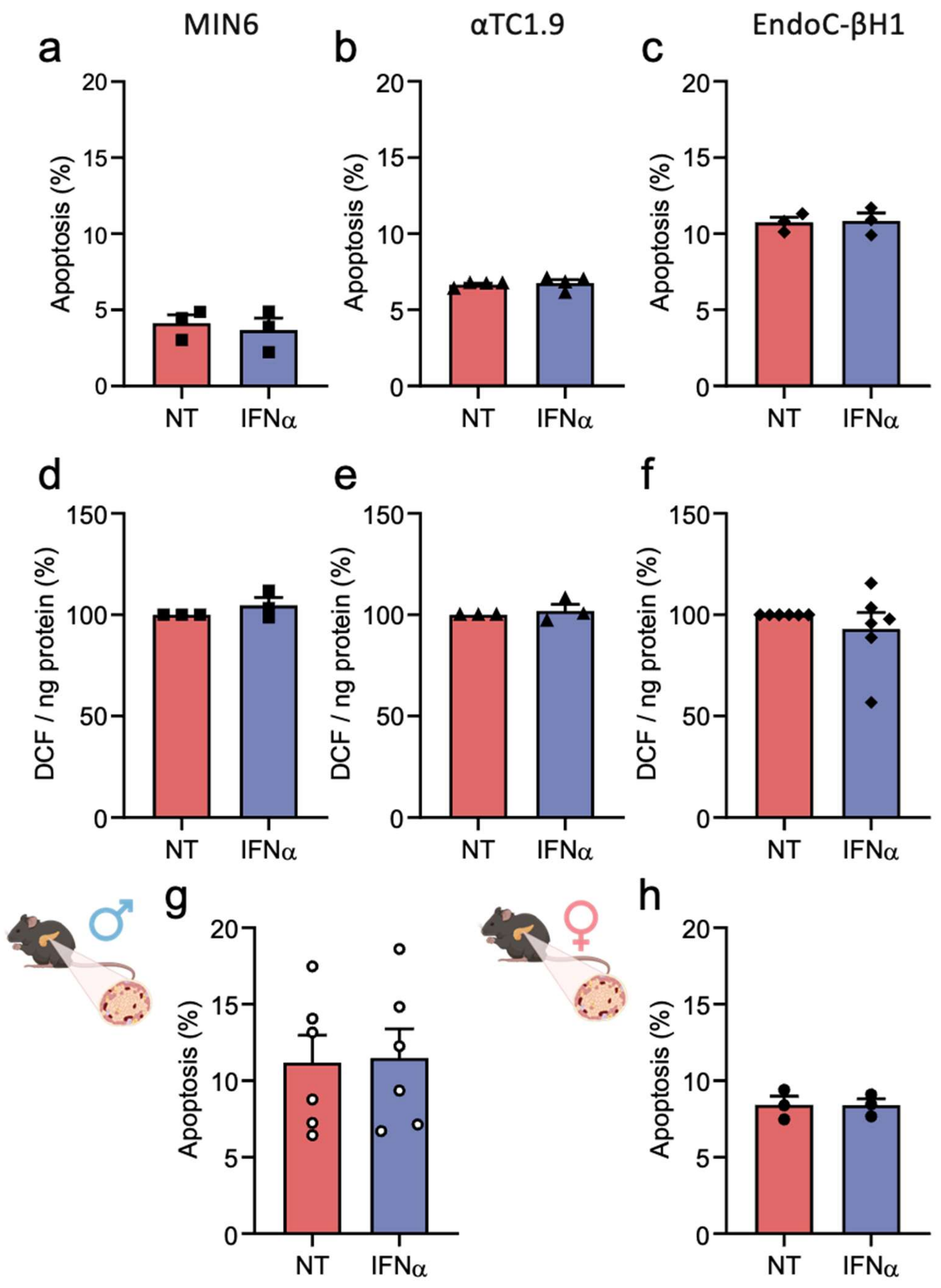


**ESM Fig. 1. IFNα does not affect cell viability or intracellular ROS production in beta and alpha cell models.**

MIN6 (**a, d**; squares), αTC1.9 (**b, e**; triangles) and EndoC-βH1 (**c, f**; diamonds) cells, and dispersed islet cells from male (**g**; open circles) and female (**h**; closed circles) GCG-Venus mice, were left untreated (NT; red) or treated with IFNα (1000 U/ml; blue) for 24 h. Apoptosis was assessed by Hoechst 33342/propidium iodide staining (**a-c, g, h**), and intracellular ROS production was assessed by DCF fluorescence (**d-f**). Results are means ± SEM of 3-6 independent experiments.

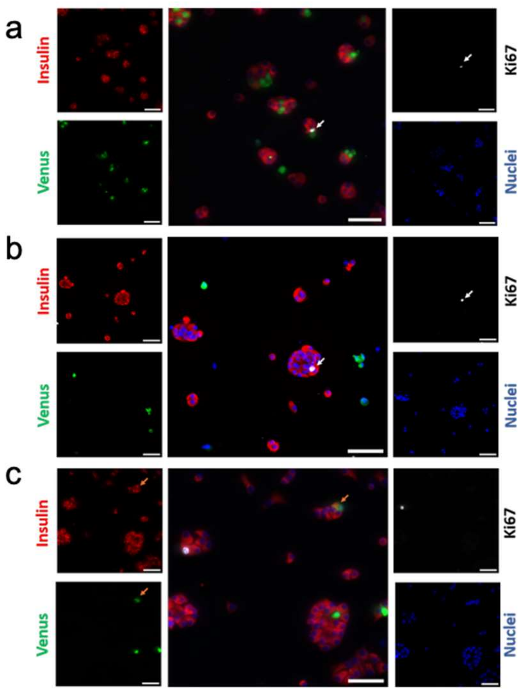


d

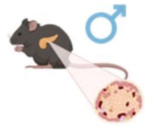

| Cell proliferation (%) | NT (*n*=4) | IFNα (*n*=4) | *p* value |
|---|---|---|---|
| Insulin⁺ | 1.17 ± 0.66 | 0.83 ± 0.43 | 0.30 |
| Venus⁺ | 0.22 ± 0.07 | 0.13 ± 0.05 | 0.45 |

| Cell population (%) | NT (*n*=4) | IFNα (*n*=4) | *p* value |
|---|---|---|---|
| Insulin⁺ | 72.36 ± 7.13 | 72.41 ± 7.44 | 0.96 |
| Venus⁺ | 14.19 ± 4.96 | 14.34 ± 4.80 | 0.64 |
| Insulin⁺/Venus⁺ | 0.92 ± 0.43 | 0.75 ± 0.23 | 0.48 |

e

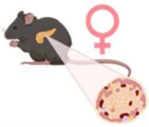

| Cell proliferation (%) | NT (*n*=4) | IFNα (*n*=4) | *p* value |
|---|---|---|---|
| Insulin⁺ | 0.47 ± 0.25 | 0.80 ± 0.07 | 0.50 |
| Venus⁺ | 0.07 ± 0.06 | 0.35 ± 0.27 | 0.43 |

| Cell population (%) | NT (*n*=4) | IFNα (*n*=4) | *p* value |
|---|---|---|---|
| Insulin⁺ | 60.50 ± 5.39 | 62.03 ± 6.64 | 0.36 |
| Venus⁺ | 22.77 ± 5.93 | 23.47 ± 6.29 | 0.37 |
| Insulin⁺/Venus⁺ | 3.52 ± 1.74 | 3.55 ± 1.44 | 0.96 |

**ESM Fig. 2. IFNα does not alter alpha or beta cell proliferation or endocrine cell proportions.**

Dispersed primary islet cells from GCG-Venus mice were left untreated (NT) or treated with IFNα (1000 U/ml) for 24 h. Alpha cells were identified by Venus fluorescence (green), beta cells by insulin immunostaining (red), proliferating cells by Ki-67 immunostaining (white), and nuclei by Hoechst 33342 staining (blue). (**a-c**) Representative single-channel and merged fluorescence images. White arrows indicate a proliferating alpha cell (**a**) and a proliferating beta cell (**b**). Orange arrows indicate cells positive for both insulin and Venus (**c**). Scale bars, 50 μm. (**d, e**) Percentages of proliferating alpha and beta cells and relative proportions of insulin-positive, Venus-positive and insulin/Venus double-positive cells in dispersed islet cultures from male (**d**) and female (**e**) mice. Results are means ± SEM from four male (116 images and 30,994 cells from NT-treated and 161 images and 35,016 cells from IFNα-treated mice analysed) and four female (98 images and 15,591 cells from NT-treated and 96 images and 14,570 cells from IFNα-treated mice analysed) mice. Paired Student's *t* test.

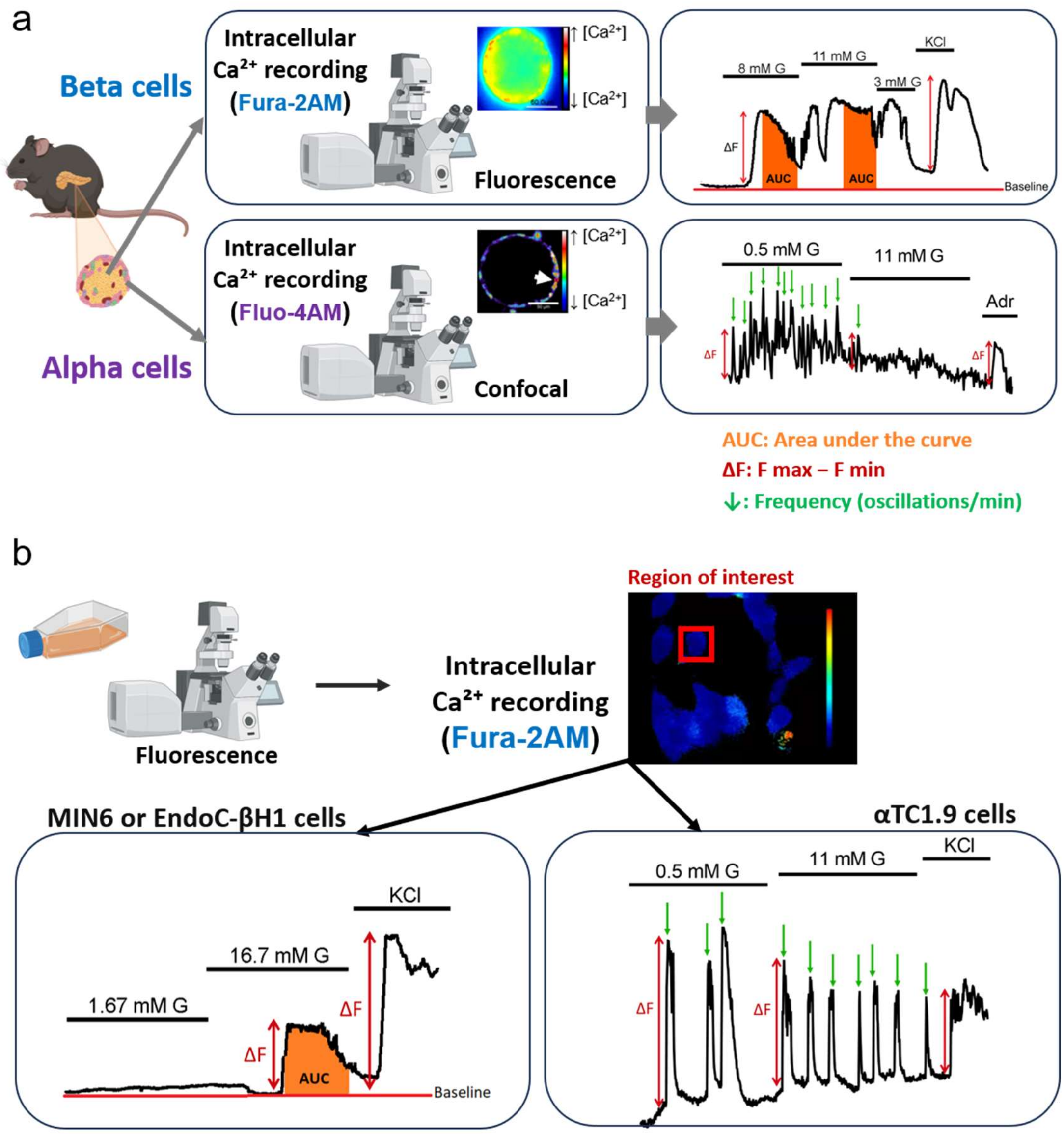


**ESM Fig. 3. Schematic representation of intracellular $Ca^{2+}$ recordings and quantified parameters in pancreatic islets and endocrine cell models.**

(**a**) Intracellular $Ca^{2+}$ dynamics were assessed in intact pancreatic islets using the ratiometric indicator Fura-2AM and in primary alpha cells within intact islets using Fluo-4AM confocal imaging. Representative traces illustrate the sequential glucose and depolarising stimuli and the parameters quantified, including area under the curve (AUC), peak amplitude (ΔF) and oscillation frequency. (**b**) Intracellular $Ca^{2+}$ dynamics were assessed using Fura-2AM in MIN6, EndoC-βH1 and αTC1.9 cells. Regions of interest were defined around individual cells. Representative traces illustrate the glucose and KCl stimulation protocols and the corresponding quantification of AUC, peak amplitude and oscillation frequency. G: glucose.

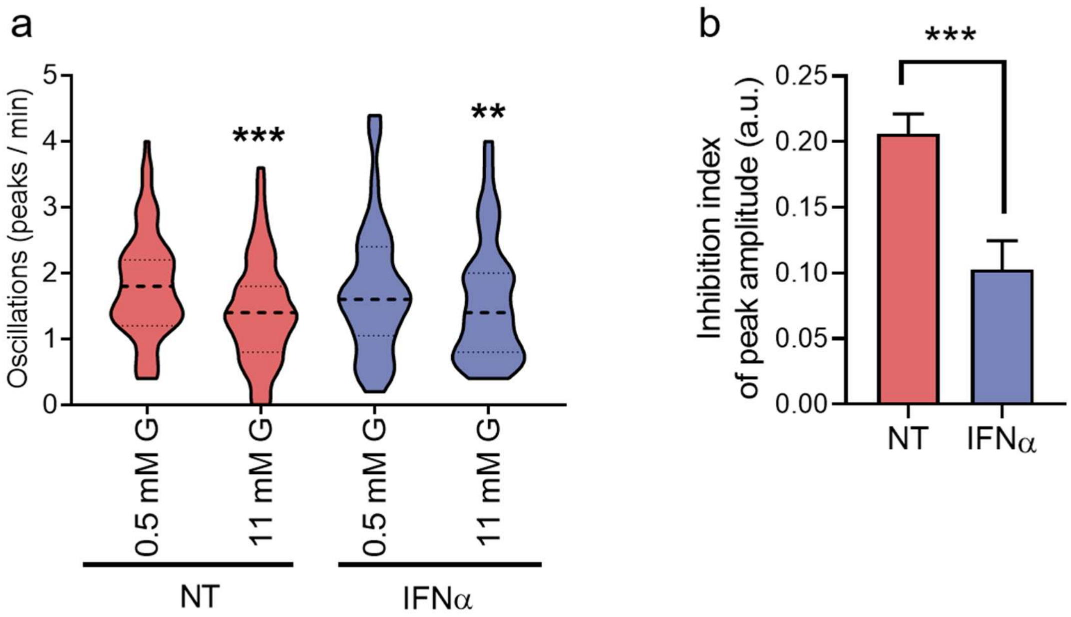


**ESM Fig. 4. Effect of IFNα on alphaTC1.9 intracellular $Ca^{2+}$ oscillations.**

αTC1.9 cells were left untreated (NT; red) or treated with IFNα (1000 U/ml; blue) for 24 h. (**a**) $Ca^{2+}$ oscillation frequency during the final 5 min of stimulation at each glucose concentration. Violin plots show the distribution, median (solid lines) and quartiles (dashed lines) from 80-87 individual registers (NT: *n*=10 recordings; IFNα: *n*=7 recordings) performed on four separate days. **$p \leq 0.01$ and ***$p \leq 0.001$ vs 0.5 mmol/l glucose within each condition; two-way ANOVA followed by Sidak's test for multiple comparisons. (**b**) Glucose inhibition index calculated from $Ca^{2+}$ peak amplitude at 11 mmol/l relative to 0.5 mmol/l glucose. Results are means ± SEM from 73-83 individual registers (NT: *n*=10 recordings; IFNα: *n*=7 recordings) performed on four separate days. ***$p \leq 0.001$ vs NT; unpaired Student's *t* test.

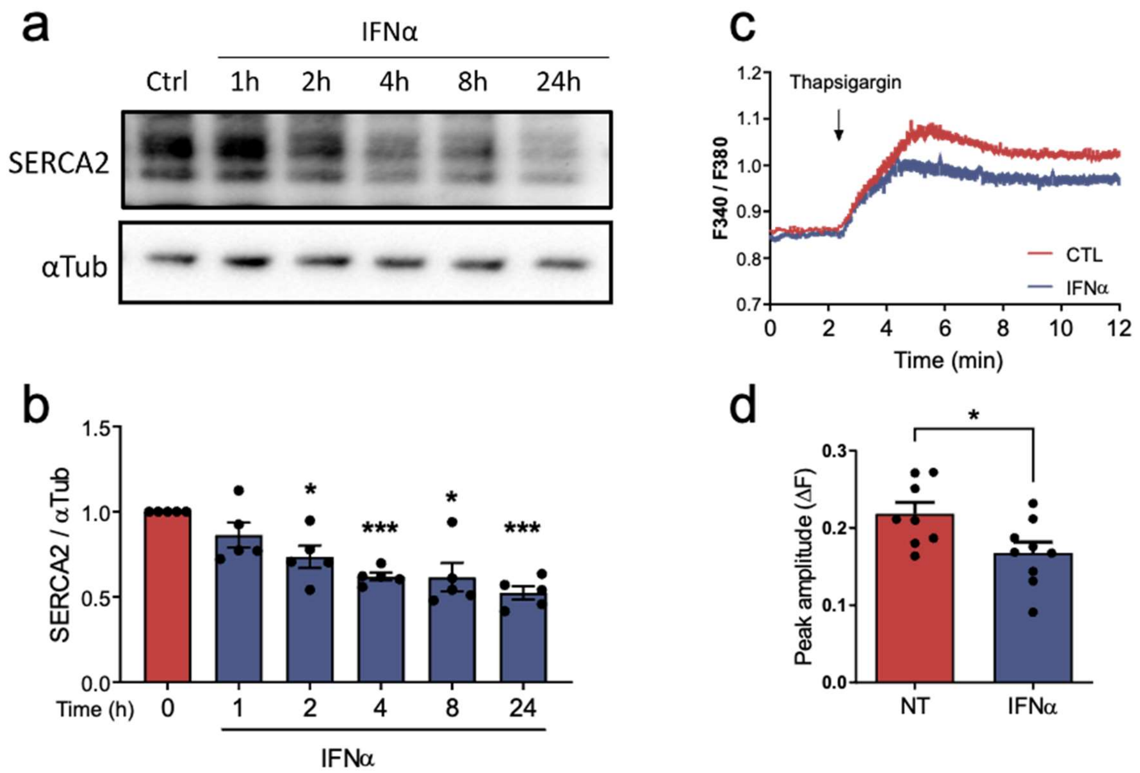


**ESM Fig. 5. IFNα reduces SERCA2 abundance and thapsigargin-induced ER $Ca^{2+}$ mobilisation in EndoC-βH1 cells.**

EndoC-βH1 cells were left untreated (NT; red) or treated with IFNα (1000 U/ml; blue) for the indicated periods (**a, b**) or for 24 h (**c, d**). (**a**) Representative immunoblot of SERCA2 abundance after 0, 1, 2, 4, 8 and 24 h of IFNα exposure, with α-tubulin (αTub) used as loading control. (**b**) Quantification of SERCA2 expression. (**c**) Representative intracellular $Ca^{2+}$ recordings following thapsigargin-induced depletion of ER $Ca^{2+}$ stores. (**d**) AUC of the thapsigargin-induced $Ca^{2+}$ response. Results are means ± SEM of 5-9 independent experiments. *$p \leq 0.05$ and ***$p \leq 0.001$ vs NT; unpaired Student's *t* test or one-way ANOVA followed by Dunnett's test for multiple comparisons, as appropriate.

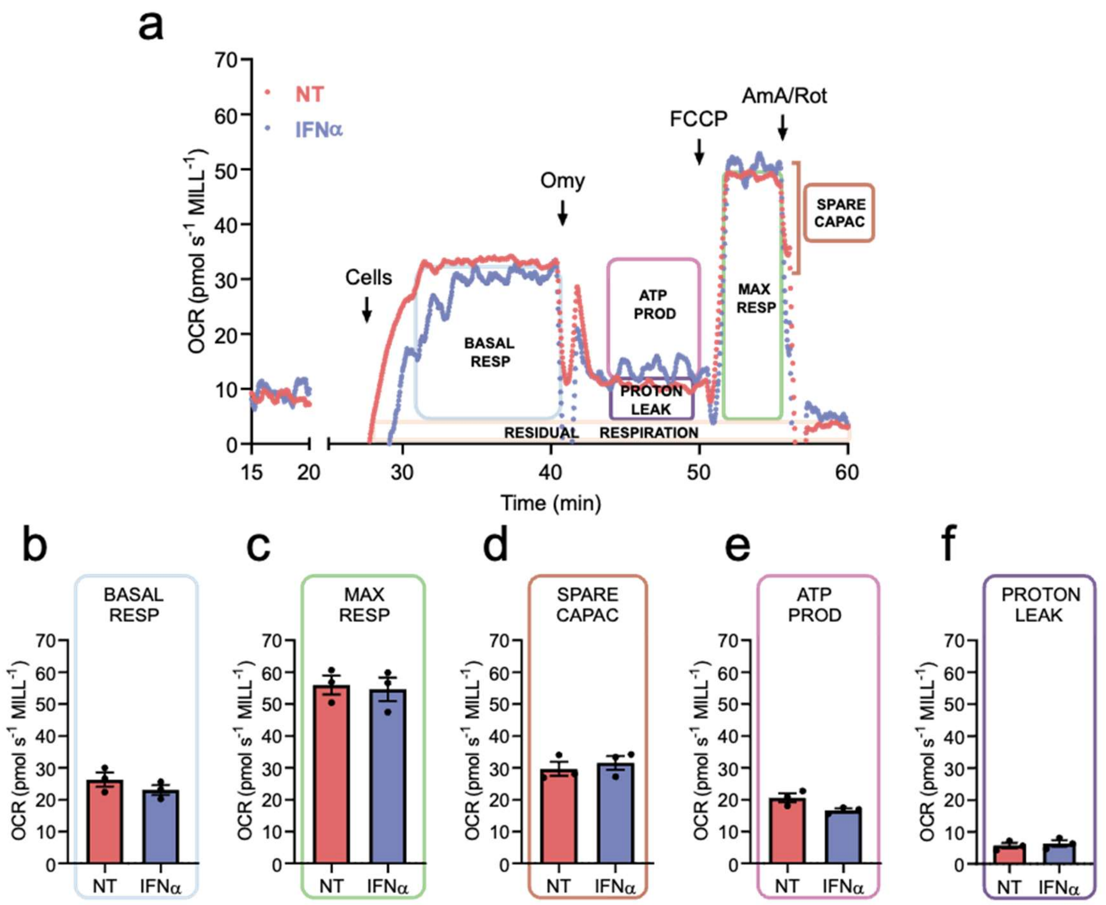


**ESM Fig. 6. IFNα does not alter mitochondrial respiration in EndoC-βH1 cells.**

EndoC-βH1 cells were left untreated (NT; red) or treated with IFNα (1000 U/ml; blue) for 24 h. (a) Representative OCR traces from untreated and IFNα-treated cells. Boxes indicate basal respiration, ATP-linked respiration, proton leak, maximal respiration, spare respiratory capacity and residual oxygen consumption. Arrows indicate additions to the chamber. (b–f) Basal respiration (b), maximal respiration (c), spare respiratory capacity (d), ATP-linked respiration (e) and proton leak (f). Results are means ± SEM of three independent experiments.

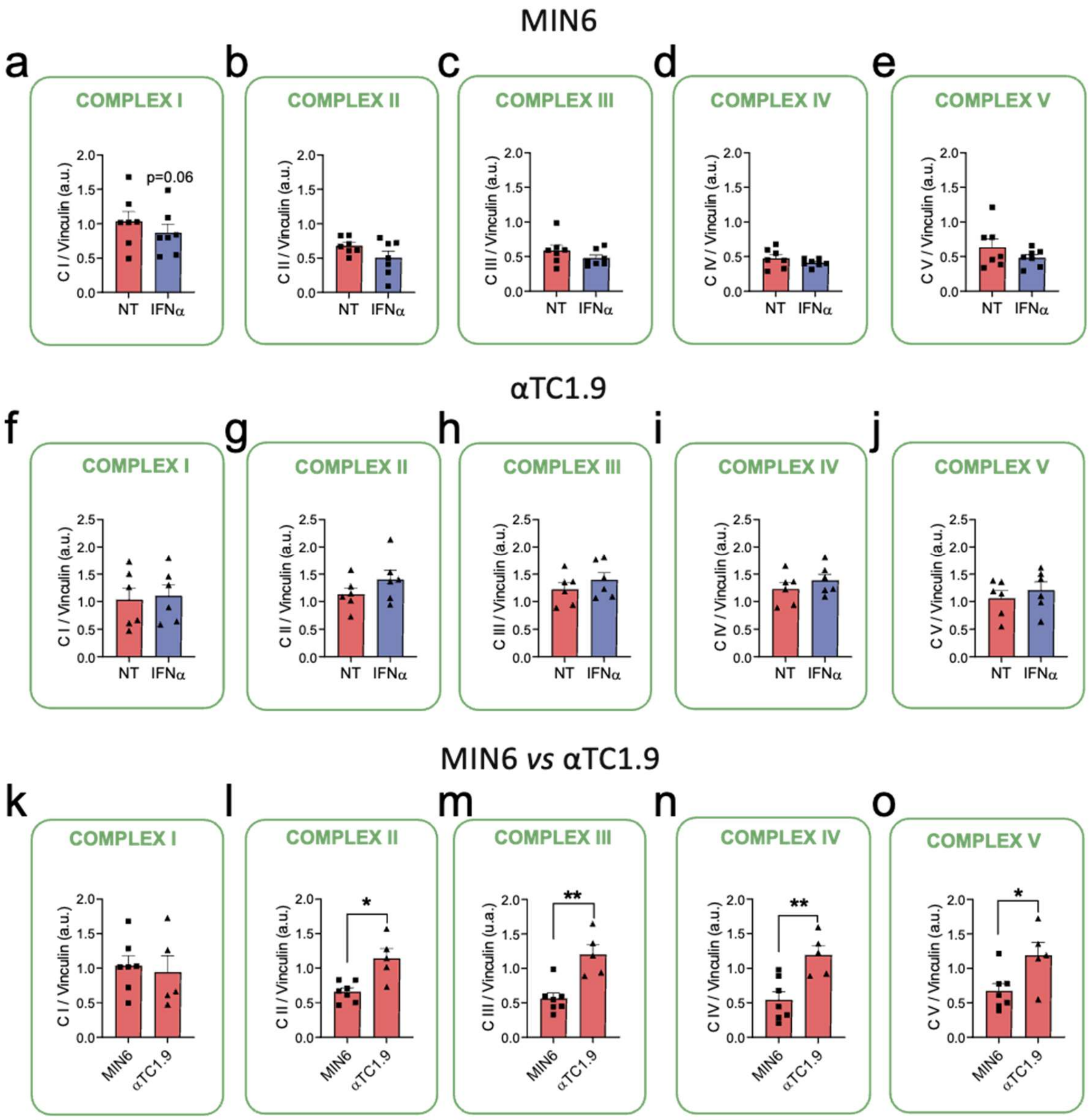


**ESM Fig. 7. IFNα does not alter mitochondrial respiratory-chain complex abundance in MIN6 or αTC1.9 cells.**

MIN6 (**a-e, k-o**; squares) and αTC1.9 (**f-o**; triangles) cells were left untreated (NT; red) or treated with IFNα (1000 U/ml; blue) for 24 h. (**a-e**) Respiratory-chain complex abundance in MIN6 cells; (**f-j**) abundance in αTC1.9 cells; and (**k-o**) comparison between the two cell lines. Values were normalised to vinculin as a total-protein loading control. Results are means ± SEM of 5-7 independent experiments. *$p \leq 0.05$ and **$p \leq 0.01$ as indicated by bars; unpaired Student's t test.

**ESM Reference list**


1. Al-Abdulla R, Ferrero H, Soriano S, Boronat-Belda T, Alonso-Magdalena P (2022) Screening of Relevant Metabolism-Disrupting Chemicals on Pancreatic β-Cells: Evaluation of Murine and Human In Vitro Models. Int J Mol Sci 23(8):4182. https://doi.org/10.3390/ijms23084182
2. Dos Santos RS, Babiloni-Chust I, Marroqui L, Nadal A (2023) Screening of Metabolism-Disrupting Chemicals on Pancreatic α-Cells Using In Vitro Methods. Int J Mol Sci 24(1):231. https://doi.org/10.3390/ijms24010231
3. Ravassard P, Hazhouz Y, Pechberty S, Bricout-Neveu E, Armanet M, Czernichow P, Scharfmann R (2011) A genetically engineered human pancreatic β cell line exhibiting glucose-inducible insulin secretion. J Clin Invest 121(9):3589–97. https://doi.org/10.1172/JCI58447
4. Dos Santos RS, Medina-Gali RM, Babiloni-Chust I, Marroqui L, Nadal A (2022) In Vitro Assays to Identify Metabolism-Disrupting Chemicals with Diabetogenic Activity in a Human Pancreatic β-Cell Model. Int J Mol Sci 23(9):5040. https://doi.org/10.3390/ijms23095040
5. Babiloni-Chust I, dos Santos RS, Medina-Gali RM, Perez-Serna AA, Encinar J-A, Martinez-Pinna J, Gustafsson J-A, Marroqui L, Nadal A (2022) G protein-coupled estrogen receptor activation by bisphenol-A disrupts the protection from apoptosis conferred by the estrogen receptors ERα and ERβ in pancreatic beta cells. Environ Int 164:107250. https://doi.org/10.1016/j.envint.2022.107250
6. Marroqui L, Dos Santos RS, Fløyel T, Grieco FA, Santin I, Op de beeck A, Marselli L, Marchetti P, Pociot F, Eizirik DL (2015) TYK2, a Candidate Gene for Type 1 Diabetes, Modulates Apoptosis and the Innate Immune Response in Human Pancreatic β-Cells. Diabetes 64(11):3808–3817. https://doi.org/10.2337/db15-0362
7. Perez-Serna AA, Dos Santos RS, Ripoll C, Nadal A, Eizirik DL, Marroqui L (2023) BCL-XL Overexpression Protects Pancreatic β-Cells against Cytokine- and Palmitate-Induced Apoptosis. Int J Mol Sci 24(6):5657. https://doi.org/10.3390/ijms24065657
8. Dos Santos RS, Marroqui L, Velayos T, Olazagoitia-Garmendia A, Jauregi-Miguel A, Castellanos-Rubio A, Eizirik DL, Castaño L, Santin I (2019) DEXI, a candidate gene for type 1 diabetes, modulates rat and human pancreatic beta cell inflammation via regulation of the type I IFN/STAT signalling pathway. Diabetologia 62(3):459–472. https://doi.org/10.1007/s00125-018-4782-0
9. Santin I, Dos Santos RS, Eizirik DL (2016) Pancreatic Beta Cell Survival and Signaling Pathways: Effects of Type 1 Diabetes-Associated Genetic Variants. In: Gillespie KM (ed) Type-1 Diabetes: Methods and Protocols. Springer, New York, NY, pp 21–54
10. Villar-Pazos S, Martinez-Pinna J, Castellano-Muñoz M, Alonso-Magdalena P, Marroqui L, Quesada I, Gustafsson J-A, Nadal A (2017) Molecular mechanisms involved in the non-monotonic effect of bisphenol-a on Ca2+ entry in mouse pancreatic β-cells. Sci Rep 7:11770. https://doi.org/10.1038/s41598-017-11995-3
11. Nadal A, Quesada I, Soria B (1999) Homologous and heterologous asynchronicity between identified α-, β- and δ-cells within intact islets of Langerhans in the mouse. J Physiol 517(1):85–93. https://doi.org/10.1111/j.1469-7793.1999.0085z.x
12. Tuduri E, Marroqui L, Soriano S, Ropero AB, Batista TM, Piquer S, Lopez-Boado MA, Carneiro EM, Gomis R, Nadal A, Quesada I (2009) Inhibitory Effects of Leptin on Pancreatic α-Cell Function. Diabetes 58(7):1616–1624. https://doi.org/10.2337/db08-1787
13. Hamilton A, Zhang Q, Salehi A, Willems M, Knudsen JG, Ringgaard AK, Chapman CE, Gonzalez-Alvarez A, Surdo NC, Zaccolo M, Basco D, Johnson PRV, Ramracheya R, Rutter GA, Galione A, Rorsman P, Tarasov AI (2018) Adrenaline Stimulates Glucagon

Secretion by Tpc2-Dependent Ca2+ Mobilization From Acidic Stores in Pancreatic α-Cells. Diabetes 67(6):1128–1139. https://doi.org/10.2337/db17-1102

14. Quesada I, Nadal A, Soria B (1999) Different effects of tolbutamide and diazoxide in alpha, beta-, and delta-cells within intact islets of Langerhans. Diabetes 48(12):2390–2397. https://doi.org/10.2337/diabetes.48.12.2390
15. Marroqui L, Martinez-Pinna J, Castellano-Muñoz M, dos Santos RS, Medina-Gali RM, Soriano S, Quesada I, Gustafsson JA, Encinar JA, Nadal A (2021) Bisphenol-S and Bisphenol-F alter mouse pancreatic β-cell ion channel expression and activity and insulin release through an estrogen receptor ERβ mediated pathway. Chemosphere 265:129051. https://doi.org/10.1016/j.chemosphere.2020.129051
16. Cunha DA, Hekerman P, Ladrière L, Bazarra-Castro A, Ortis F, Wakeham MC, Moore F, Rasschaert J, Cardozo AK, Bellomo E, Overbergh L, Mathieu C, Lupi R, Hai T, Herchuelz A, Marchetti P, Rutter GA, Eizirik DL, Cnop M (2008) Initiation and execution of lipotoxic ER stress in pancreatic β-cells. J Cell Sci 121(14):2308–2318. https://doi.org/10.1242/jcs.026062
